# Colocalization of blood cell traits GWAS associations and variation in PU.1 genomic occupancy prioritizes causal noncoding regulatory variants

**DOI:** 10.1101/2023.03.29.534582

**Authors:** Raehoon Jeong, Martha L. Bulyk

**Affiliations:** Division of Genetics, Department of Medicine, Brigham and Women’s Hospital and Harvard Medical School, Boston, MA 02115, USA; Bioinformatics and Integrative Genomics Graduate Program, Harvard University, Cambridge, MA 02138, USA; Department of Pathology, Brigham and Women’s Hospital and Harvard Medical School, Boston, MA 02115, USA

## Abstract

Genome-wide association studies (GWAS) have uncovered numerous trait-associated loci across the human genome, most of which are located in noncoding regions, making interpretations difficult. Moreover, causal variants are hard to statistically fine-map at many loci because of widespread linkage disequilibrium. To address this challenge, we present a strategy utilizing transcription factor (TF) binding quantitative trait loci (bQTLs) for colocalization analysis to identify trait associations likely mediated by TF occupancy variation and to pinpoint likely causal variants using motif scores. We applied this approach to PU.1 bQTLs in lymphoblastoid cell lines and blood cell traits GWAS data. Colocalization analysis revealed 69 blood cell trait GWAS loci putatively driven by PU.1 occupancy variation. We nominate PU.1 motif-altering variants as the likely shared causal variants at 51 loci. Such integration of TF bQTL data with other GWAS data may reveal transcriptional regulatory mechanisms and causal noncoding variants underlying additional complex traits.

---

A recurring challenge in genome-wide association studies (GWAS) is the difficulty of identifying causal variants, as well as formulating corresponding variant-to-function (V2F) hypotheses^1^. Pinpointing causal variants is important as it guides subsequent validation experiments^2–4^ and development of potential therapies^5^. More precise identification of causal variants (*e.g.*, fine-mapping) also leads to better genetic risk predictions across various traits and diseases^6, 7^. However, widespread linkage disequilibrium (LD) typically prevents effective statistical fine-mapping, especially for common variants^1, 8^. Moreover, most of the genome-wide significant loci are noncoding and likely have regulatory functions; in practice, noncoding variants are much harder to interpret than coding variants because predicting the effects of noncoding variants on transcription factor (TF) binding *in vivo* is challenging. Since variants predicted to affect TF binding across the genome have been shown to explain a large proportion of genetic associations to traits (*i.e.*, heritability enrichment)^9, 10^, many studies have examined whether trait-associated variants overlap a TF binding site motif within the corresponding TF ChIP-seq peak^8, 11^, but data demonstrating the variants’ effects on *in vivo* TF binding are necessary to imply causality of the variant on TF binding. Therefore, an approach to effectively pinpoint regulatory variants and their effects on *in vivo* TF binding at individual GWAS loci is essential.

Previous studies have utilized expression quantitative trait loci (eQTL) or methylation QTL (mQTL) colocalization to learn about regulatory mechanisms (*e.g.*, causal genes) at GWAS loci^12–16^. Statistical colocalization specifically tests the hypothesis that genetic signals are shared between a pair of traits (*e.g.* eQTL and GWAS), whereas positional overlap of associations to two traits alone leads to many false positives^13, 17^. However, a key weakness in eQTL or mQTL colocalization analysis is the inability to pinpoint a causal regulatory variant effectively because colocalization analyses are not inherently aimed at identifying the causal variant, and LD typically prevents statistical fine-mapping at single-variant resolution^14^.

Here, we have developed a strategy 1) to analyze colocalization of TF binding QTLs (bQTLs) (*i.e.*, genomic loci where TF occupancy level, as measured by ChIP-seq, is significantly associated with a genetic variant) at GWAS loci to highlight TF binding sites that potentially mediate the GWAS associations^18^, and 2) to utilize TF motif models to nominate variants altering a motif of the corresponding TF at those binding sites as likely shared causal regulatory variants underlying both TF binding variation and the GWAS traits (Fig. 1a). TF bQTLs are fundamentally different from eQTLs and mQTLs in that TF bQTLs point to likely causal variants because they are often driven by the corresponding TF motif-altering variants^19, 20^. To our knowledge, this is the first attempt to perform TF bQTL colocalization analysis with GWAS data to fine-map putative causal variants that affect *in vivo* TF binding.

**Fig. 1:**
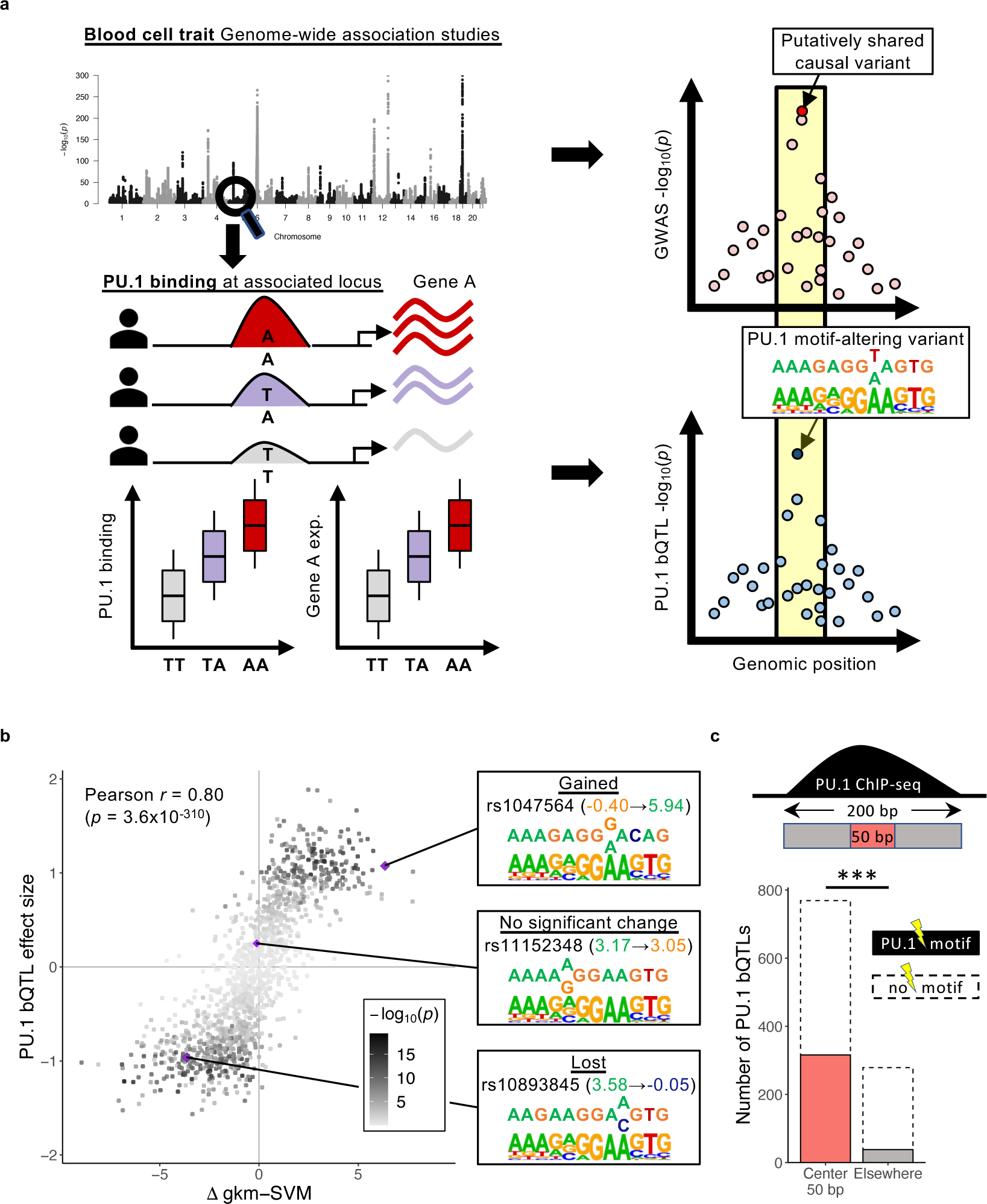
Relevance of PU.1 bQTLs in LCLs to blood cell trait associations. (**a**) (Left) Blood cell trait-associated loci may have overlapping PU.1 bQTLs and, potentially, expression QTL (eQTL) associations. (Right) Significant colocalization suggests that the causal variants are shared. If there is a PU.1 motif-altering variant at a colocalized PU.1 bQTL, the variant is likely to be the shared causal variant. (**b**) Comparison of changes in motif score (Δ gkm-SVM) and estimated bQTL effect sizes at PU.1 motif-altering variants within 200bp PU.1 ChIP-seq peaks. The color represents the –log_10_(*p*) of PU.1 bQTL association (linear regression). (**c**) Number of significant PU.1 bQTLs with PU.1 motif-altering variants at each region within the 200bp PU.1 ChIP-seq peaks. ***: *p* < 2.2×10^−16^ (Fisher’s exact test).

We carried out this strategy with blood cell trait GWAS^21^ and bQTL data for the hematopoietic master regulator PU.1 from lymphoblastoid cell lines (LCLs)^19, 22^, which are immortalized B cell lines. PU.1 bQTLs in neutrophils have been found previously to colocalize with immune disease susceptibility loci but were not used to fine-map the causal variants^18^. Blood cell traits (*e.g.*, lymphocyte counts, hemoglobin concentrations) are indicators of various diseases; for instance, individuals with low lymphocyte counts are more susceptible to infections, including severe COVID-19^23–25^. Consistent with PU.1’s role in specifying myeloid and lymphoid lineages during hematopoiesis^26, 27^ and its expression throughout progenitor cell types^28^ (Supplementary Fig. 1), a recent fine-mapping analysis of blood cell trait GWAS reported that PU.1 was the TF with the highest number of fine-mapped noncoding variants altering its DNA binding site motif^11^, suggesting that PU.1 motif-altering variants might drive many blood cell trait association signals.

In order to identify blood cell trait associations that may be driven by a variant altering PU.1 binding, we analyzed publicly available PU.1 ChIP-seq data from LCLs across 49 individuals^19, 22^ and identified 1497 PU.1 bQTLs. Next, PU.1 bQTLs colocalized with at least one blood cell trait association at 69 loci; for 51 of these loci, we identified PU.1 motif-altering variants as the likely causal variants. Thus, our approach allowed us to overcome the limitations of statistical fine-mapping in resolving these GWAS signals to single causal variants. By incorporating chromatin accessibility, histone mark, and transcriptome data for LCLs, we identified several putative causal genes for traits, including lymphocyte and monocyte counts. More broadly, our results illustrate the utility of TF bQTL datasets for fine-mapping trait-associated noncoding loci and in generating mechanistic, V2F models of gene dysregulation for traits of biomedical importance.

## Results

### PU.1 motif-altering variants are likely causal for PU.1 bQTL associations

First, we reanalyzed available PU.1 ChIP-seq data for LCLs from 49 individuals^19, 22^. These individuals are all of European ancestry, and their genotypes are available through the 1000 Genomes Project^29^ (Supplementary Table 1). After peak calling and normalization of the PU.1 ChIP-seq read counts, we tested for significant genetic associations with common variants (minor allele frequency (MAF) > 0.05) within 100 kb of each ChIP-Seq peak. In total, we identified 1497 significant PU.1 bQTLs (FDR < 5%).

We next inspected the contribution of PU.1 motif-altering variants to PU.1 bQTLs. First, we verified that PU.1-occupied regions were enriched for a match to the PU.1 binding site motif, identified by a position weight matrix (PWM), near the center of the ChIP-Seq peaks (Extended Data Fig. 1a), suggesting that most of these sites are bound directly by PU.1. Next, we evaluated whether PU.1 motif-altering variants affect PU.1 binding by training a motif score model gkm-SVM^30, 31^ to learn gapped *k*-mers that are overrepresented in PU.1-occupied sequences. The model captured both PU.1 and PU.1:IRF composite motifs (Extended Data Fig. 1b), the latter of which reflects PU.1 binding to DNA as a heterodimer with either IRF4 or IRF8^32^. Changes in gkm-SVM scores have been shown to predict effects of variants on TF binding better than PWMs^33^, which imprecisely assume each nucleotide to affect binding independently. Consistent with our expectations, the predicted change in gkm-SVM scores for single nucleotide polymorphism (SNP) within PU.1 motifs were significantly correlated with estimated PU.1 bQTL effect sizes (Pearson *r* = 0.80, *p* = 3.6×10^−310^) (Fig. 1b, Supplementary Table 2). This strong positive correlation supports the model that PU.1 motif-altering variants, if present, are likely causal for those PU.1 bQTLs. Furthermore, significant PU.1 bQTLs with a motif-altering variant (determined by gkm-SVM) showed that such variants are more concentrated towards the peak centers compared to PU.1 bQTLs without one (Fig. 1c, two-sided Fisher’s exact test *p* = 3.1×10^−18^), consistent with the expectation that PU.1 motif-altering variants directly affect PU.1 occupancy. Hence, we considered that PU.1 bQTLs colocalized with blood cell traits association would likely be driven by PU.1 motif-altering variants, if present (Fig. 1a).

### PU.1 binding sites and PU.1 bQTLs in LCLs are enriched for blood cell trait association

To verify the relevance of these PU.1 bQTLs for investigations of blood cell traits, we evaluated whether the PU.1 bQTLs are more likely to be significantly associated with each of the 28 blood cell traits (Supplementary Table 3) than expected by chance. We analyzed blood cell traits GWAS data from UK Biobank^21^. As a background expectation, we constructed 250 sets of null variants matched with PU.1 bQTL lead variants for allele frequency, number of tagging variants (LD *r*^2^ > 0.5), and distance to the closest transcription start site (TSS). The significant PU.1 bQTLs were more likely to tag lead variants associated (*i.e.*, *p* < 5×10^−8^) with myeloid lineage traits (*e.g.* monocyte and neutrophil count) and lymphoid lineage traits (*e.g.* lymphocyte count) than the sets of null variants (empirical adjusted *p* < 0.05) (Fig. 2a), which is consistent with the known role of PU.1 in myeloid and lymphoid differentiation^26, 27^. In contrast, PU.1 bQTLs were not enriched for other traits like type 2 diabetes or height (Extended Data Fig. 1c).

**Fig. 2:**
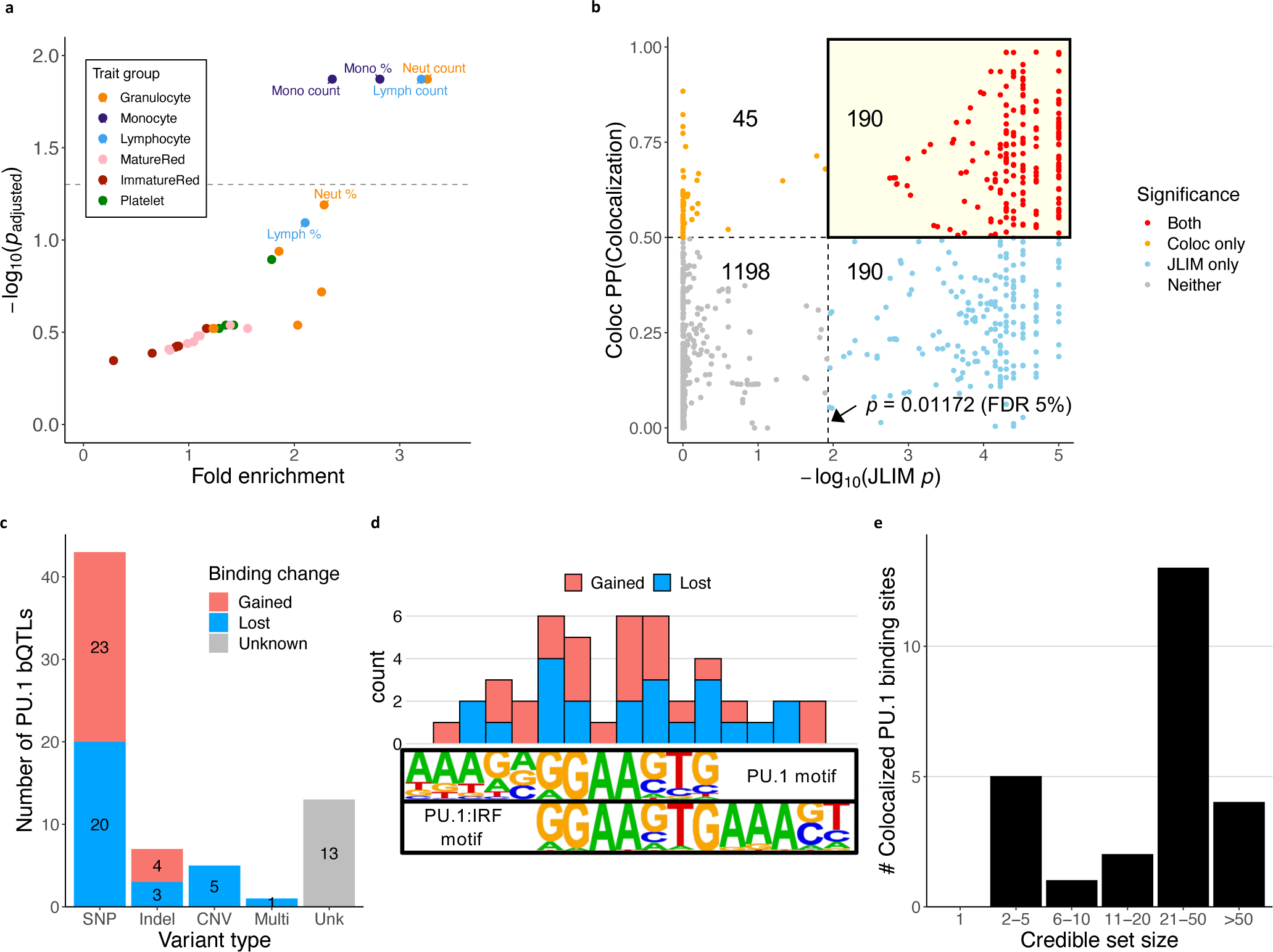
Colocalization of blood cell traits GWAS and PU.1 bQTLs. (**a**) Enrichment of PU.1 bQTLs for associations to specific blood cell traits. Traits with empirical adjusted *p* < 0.05 (above the dashed line) are labeled. Abbreviations of blood cell traits are described in Supplementary Table 3. (**b**) Colocalization results from JLIM and Coloc. Each point is a PU.1 bQTL - Trait pair. The number shown in each quadrant is the number of points within the significance category. Dashed lines indicate the respective significance thresholds (JLIM: *p* < 0.01172 (FDR 5%), Coloc: PP(colocalized) > 0.5). (**c**) The types of putative causal variants at colocalized PU.1 bQTLs that alter PU.1 motifs or the copy number of the PU.1 occupancy site. SNPs, indels, and multi-variants alter PU.1 motifs. CNV: copy number variation altering copy number of PU.1 binding sites; Multi: multiple variants in perfect LD (*r*^2^ = 1) within a PU.1 motif sequence; Unk (Unknown): No variant altering PU.1 motif sequence or its copy number. (**d**) Number of PU.1 motif-altering SNPs at each nucleotide position at colocalized PU.1 binding sites. Motif logos are from Homer database. (**e**) Blood cell trait GWAS credible set size at loci with colocalized PU.1 bQTLs and a PU.1 motif-altering variant. Only 25 loci with fine-mapping result in Vuckovic et al. 2020 are represented.

### PU.1 bQTL colocalization with blood cell trait associations

To identify candidate loci to test for potential colocalization of PU.1 bQTL and blood cell trait associations, we filtered all significant PU.1 bQTLs for loci with at least one blood cell trait association at *p* < 10^−6^. This resulted in a total of 1621 such PU.1 bQTL-trait pairs, comprising 367 unique loci. We then applied two distinct colocalization methods – JLIM^13^ and Coloc^12^ – to test for robust colocalization (Supplementary Table 4). Chun and colleagues showed with simulated data that each method can show different performance depending on the LD structure of the loci^13^; therefore, we reasoned that requiring significant colocalization by both methods would enrich true positive cases. We used a significance threshold of *p* < 0.01172 (FDR < 5%) for JLIM and posterior probability of colocalization (PP(Colocalization)) > 0.5 for Coloc.

The statistically significant colocalization of PU.1 bQTL-trait pairs identified by JLIM and Coloc were overall consistent (Pearson *r* = 0.73, *p* = 6.8×10^−270^; Fig. 2b). We identified a total of 190 (11.7%) PU.1-trait pairs, spanning 69 unique loci, that were significant by both methods (Fig. 3). Across the blood cell traits, those related to white blood cells (*e.g.* white blood cell count, lymphocyte count, neutrophil count) showed a higher proportion of the tested loci showing high-confidence colocalization than red blood cell or platelet traits (Fig. 3a), similar to the enrichment of tagging variants observed in Fig. 1b. We also found 1196 (73.8%) cases where a variant that was significant for both PU.1 bQTL and blood cell traits did not exhibit significant colocalization by either JLIM or Coloc, highlighting the importance of performing colocalization analysis to distinguish loci with statistical evidence of shared causal variants from those where the variants associated with each trait are merely in LD with each other^17^. The remaining 235 (14.5%) pairs showed discordant results between the two methods, which could potentially stem from lack of statistical power due to weak association signals or many variants showing high LD with the lead variant (Supplementary Fig. 2, Supplementary Note). This discrepancy justifies the rationale of applying both methods to identify high-confidence colocalization.

**Fig. 3:**
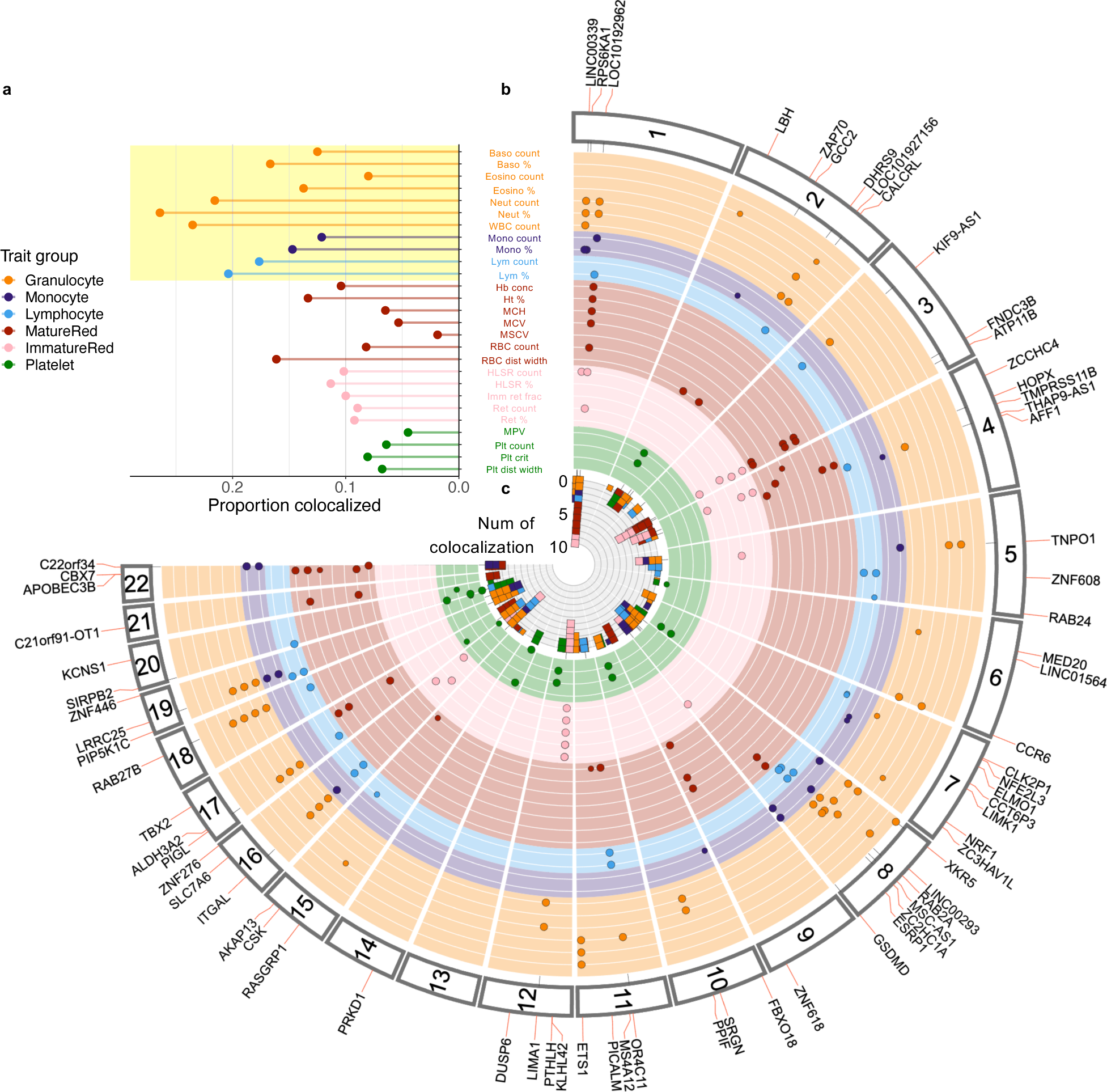
Distribution of colocalized loci across the genome. (**a**) Proportion of tested loci with significant colocalization. The colors represent the trait groups. The blood cell traits highlighted in yellow correspond to white blood cell traits. Abbreviations of blood cell traits are described in Supplementary Table 3. (**b**) Fuji plot depicting the genomic distribution of blood cell trait-associated loci that show high-confidence colocalization with PU.1 bQTLs. The colors are as in panel a. (**c**) The stacked bar plot at the center shows the number of traits each PU.1 bQTL colocalizes with.

Most (56/69) loci showing high-confidence colocalization had some biologically plausible putative causal variants (*i.e.*, directly affecting a PU.1 binding sequence) (Fig. 2c, Extended Data Fig. 2a, Supplementary Table 5). 43 (62.3%) loci had a SNP altering a PU.1 motif, while 7 (10.1%) had a short insertion or deletion (indel) variant. In addition, there was one locus where two adjacent SNPs were in perfect LD (*r*^2^= 1) and altered a single PU.1 motif sequence (Extended Data Fig. 2a and Supplementary Table 6). These SNPs and short indels showed a balance of gained and lost PU.1 binding (two-sided binomial test *p* = 0.67), and changes in gkm-SVM motif scores were highly correlated with the estimated PU.1 bQTL effect sizes (Pearson *r* = 0.89, *p* = 5.2×10^−18^) (Extended Data Fig. 2b). The PU.1 motif-altering SNPs at colocalized loci are distributed within the PU.1 or PU.1:IRF motif, with the highest frequencies at the core “GGAAG” positions (Fig. 2d and Supplementary Table 7). We retrieved fine-mapping results for 25 colocalized loci with a PU.1 motif-altering variant (*i.e.* SNP or indel) from a recent blood cell trait GWAS study^8^ (Supplementary Note). 19 of these 25 (76%) loci had more than 10 variants in the 95% credible set (*i.e.*, minimal set of variants that have 95% posterior probability of containing the causal variant), none of which was fine-mapped to a single variant (Fig. 2e and Supplementary Table 8). Despite difficulty in fine-mapping due to LD structure, we were able to pinpoint putative causal variants in these loci using a specific TF’s (*i.e.*, PU.1) motif information. There were also 5 loci with large deletions that completely removed the PU.1 binding site, which we were able to uncover because the 1000 Genomes Project (1KGP)^29^ genotypes included structural variants (Extended Data Fig. 2c); whether the deletions are true causal variants will need to be tested experimentally in future studies.

Pinpointing likely causal regulatory variants allowed us to derive specific hypotheses about gene regulatory mechanisms that are perturbed by the variants, as described below. We show one example where a PU.1 motif-altering SNP (rs12517864) represents a secondary expression QTL (eQTL) (*i.e.*, a weaker signal independent from the strongest, primary eQTL) to *ZNF608* in LCLs, and only this secondary signal colocalizes with lymphocyte count association (Fig. 4); an eQTL-centric analysis in LCLs would have missed this locus without accounting for multiple independent signals, highlighting the power of the use of TF bQTL data in colocalization analysis with GWAS data. Two other examples show reporter assay results corroborating the regulatory effects of PU.1 motif-altering variants identified in colocalized loci (Fig. 5 and 6).

**Fig. 4:**
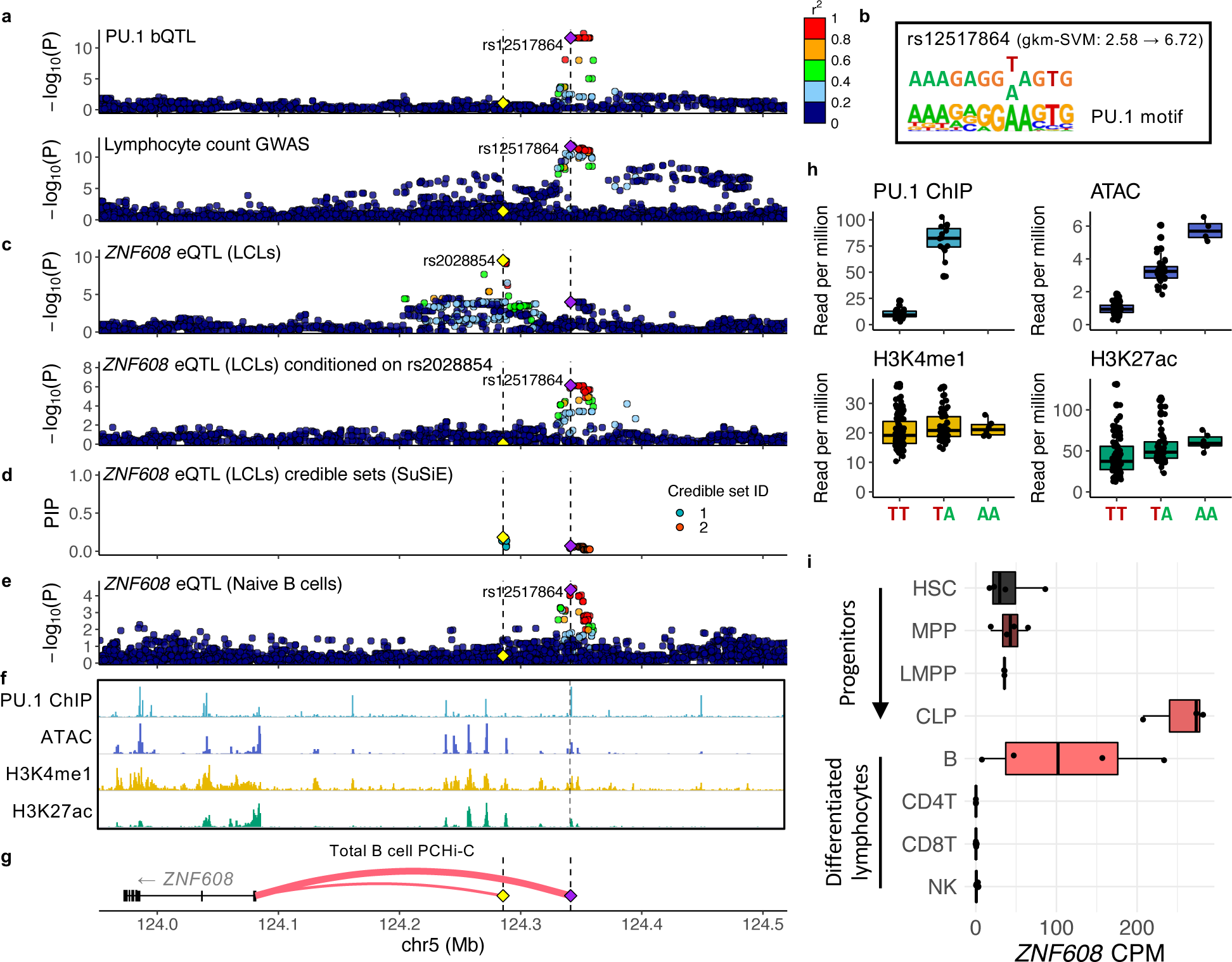
PU.1 motif alteration pinpoints a lymphocyte count-associated variant that is a secondary *ZNF608* eQTL variant. (**a, c-e, g**) PU.1 motif-altering variant rs12517864 is shown as a purple diamond, and the *ZNF608* eQTL lead variant rs2028854 is shown as a yellow diamond. Vertical dashed lines mark the position of these two variants. Unless noted otherwise, points are colored by LD *r*^2^ with respect to rs12517864. (**a**) PU.1 bQTL and lymphocyte count association signals. (**b**) The effect of rs2028854 on the sequence with respect to the PU.1 binding motif. (**c**) (Top) Primary *ZNF608* eQTL signals in LCLs. LD *r*^2^ is calculated with respect to rs2028854, the lead variant. (Bottom) *ZNF608* eQTL signals in LCLs conditioned on the rs2028854 dosage. (**d**) Fine-mapping result of *ZNF608* eQTL signals in LCLs, using SuSiE. Points are colored by the credible set they belong to. PIP: Posterior inclusion probability. (**e**) *ZNF608* eQTL association signals in naïve B cells (DICE). (**f**) Genome tracks of PU.1 ChIP-seq, ATAC-seq, H3K4me1 and H3K27ac ChIP-seq assayed in GM12878. (**g**) Gene track showing *ZNF608* and the two variants. The weights of the red curves indicate the CHiCAGO scores calculated in Javierre et al. 2016. (**h-i**) On top of the box plots, all the data points are shown. (**h**) The effect of rs12517864 dosage on various molecular phenotypes shown in panel f. For PU.1 ChIP-seq data, there weren’t any individuals with homozygous alternate allele (AA). (**i**) *ZNF608* expression levels (count per million) through lymphocyte differentiation and across various lymphocyte types. HSC: hematopoietic stem cell, MPP: multipotent progenitor, LMPP: lymphoid-primed multipotent progenitor, CLP: common lymphoid progenitor, B: B cell, CD4T: CD4^+^ T cell, CD8T: CD8^+^ T cell, NK: natural killer cell.

**Fig. 5:**
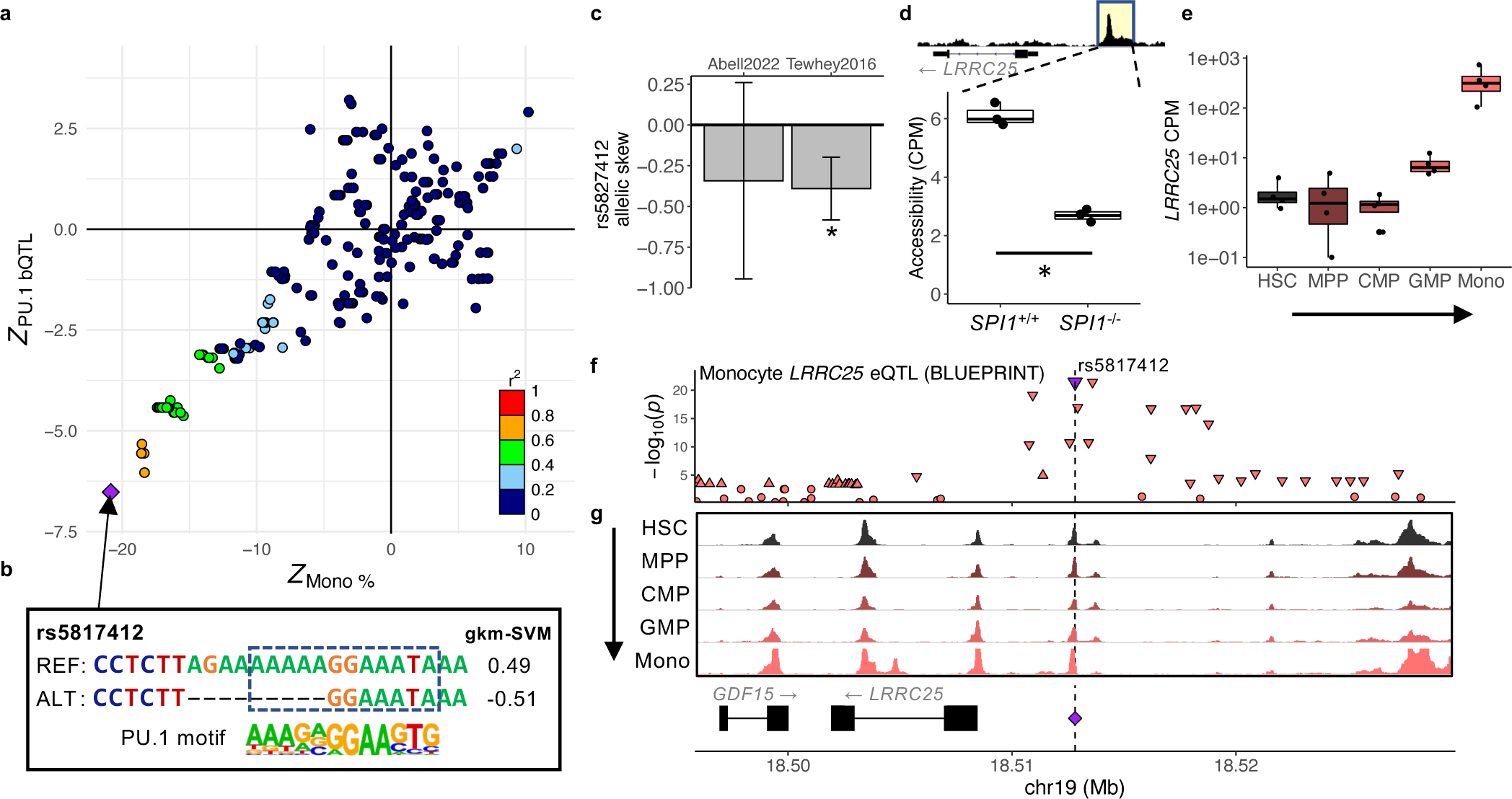
PU.1 motif-altering deletion rs5827412 at *LRRC25* locus associated with lower monocyte counts. (**a**) PU.1 bQTL and monocyte percentage association signals colocalize. (**b**) The effect of rs5827412 on the PU.1 motif. (**c**) Reduced reporter activity by rs5827412 in log2 fold change. Error bars indicate 95% confidence intervals. *: adjusted *p* < 0.05. (**d-e**) Boxplots are formatted as in Fig 4. (**d**) A boxplot showing PU.1-dependent reduction in chromatin accessibility levels (count per million) at the regulatory element surrounding rs5827412 in control pro-B cell lines (*SPI1*^+/+^) and counterparts with *SPI1* knocked out (*SPI1*^−/-^). Regions highlighted in yellow marks the accessible region corresponding to the boxplot. *n* = 3 for each condition. *: DESeq2 adjusted *p* < 0.05. (**e**) A boxplot showing *LRRC25* expression levels (count per million) through monocyte differentiation. HSC: hematopoietic stem cell, MPP: multipotent progenitor, CMP: common myeloid progenitor, GMP: granulocyte-macrophage progenitor, Mono: monocyte. (**f-g**) Purple triangle and diamond, as well as the dashed line, mark rs5827412. (**f**) Monocyte *LRRC25* eQTL association. Downward and upward triangles indicate the direction of effect (down- and up-regulation, respectively) for variants with *p* < 1×10^−3^. (**g**) ATAC-seq tracks as fold enrichment over average (range 0-40) for various blood cell types through monocyte differentiation.

**Fig. 6:**
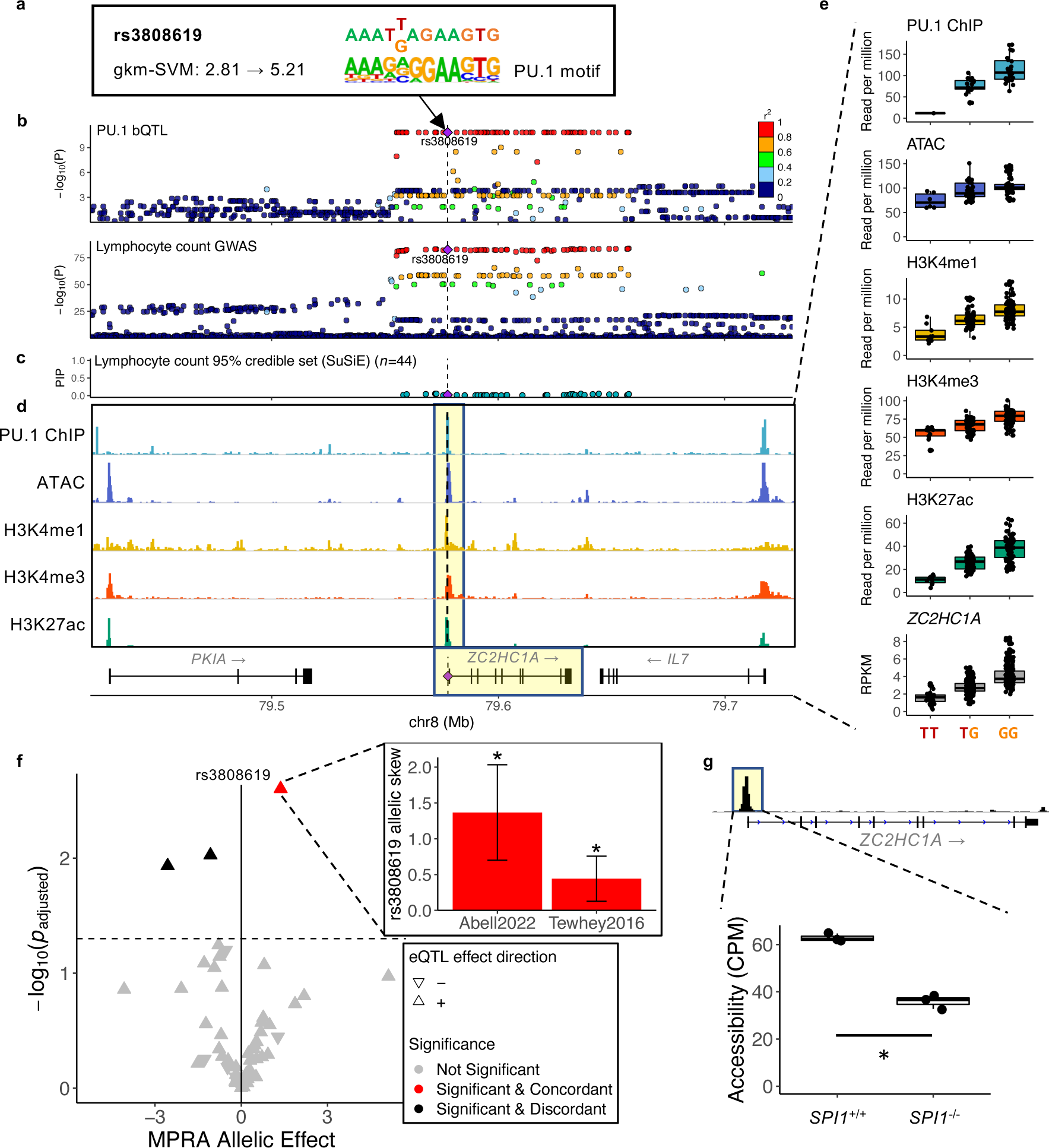
*ZC2HC1A* locus: PU.1 motif-alteration highlights a regulatory variant among those in high LD. (**a-d**) PU.1 motif-altering variant rs3808619 is shown as a purple diamond. Vertical dashed line also mark the position of this variant. (**a**) The effect of rs3808619 on the PU.1 composite motif. (**b**) PU.1 bQTL and lymphocyte count association signal at the *ZC2HC1A* locus. (**c**) Posterior inclusion probability (PIP) of variants in the 95% credible set of lymphocyte count association at the *ZC2HC1A* locus. (**d**) Genome tracks of PU.1 ChIP-seq, ATAC-seq, H3K4me1, H3K4me3, H3K27ac ChIP-seq assayed in GM12878. The highlighted regions correspond to molecular phenotypes with QTL associations in **e**. (**e**) The effect of rs3808619 dosage on various molecular phenotypes shown in panel d. Box plots are formatted as in Fig. 4. (**f**) Regulatory effects of rs3808619 and 58 tagging variants in a reporter assay. MPRA allelic effect corresponds to log2 fold change of regulatory activity of the oligo sequence with the alternate allele over that with the reference allele. The inset shows the allelic skew estimates with 95% confidence intervals from Abell et al. and Tewhey et al. *: adjusted *p* < 0.05. (**g**) PU.1-dependent reduction in chromatin accessibility levels (count per million) at the regulatory element surrounding rs3808619 in control pro-B cell lines (*SPI1*^+/+^) and counterparts with *SPI1* knocked out (*SPI1*^−/-^). *n* = 3 for each condition. *: DESeq2 adjusted *p* < 0.05. The panel is formatted as in Fig. 5d.

### bQTL colocalization reveals a putative causal variant that is not the primary eQTL

Causal genes at a trait-associated locus frequently have been identified using eQTL data for nearby genes^14, 34^. However, eQTLs can often have multiple independent signals^14^, and these signals detected in any one cell type may not all be associated with a GWAS trait, such as if the regulatory effects manifest themselves only in certain cellular contexts. This complicates colocalization analyses that often assume a single shared causal variant at a locus^12, 13^. In contrast, TF bQTLs capture regulatory effects of individual regulatory elements. Therefore, TF bQTL colocalization analysis can isolate the effects of variants on specific regulatory elements, lowering the probability of multiple causal variants compared to that of eQTLs.

For example, the *ZNF608* locus shows significant colocalization of PU.1 bQTL and lymphocyte count association (Fig. 4a, Extended Data Fig. 3a). Although the molecular function of *ZNF608* remains unclear, a study of follicular lymphoma (FL), a type of cancer in which B lymphocytes divide uncontrollably, found this gene to be among the 39 genes significantly enriched for missense or predicted-loss-of-function (pLOF) somatic mutations in FL patients^35^, suggesting it plays a role in B lymphocyte development. The associated PU.1 binding site is located about 257 kilobases (kb) upstream of the *ZNF608* promoter, and the SNP rs12517864 that increases the PU.1 binding motif score (0.68→2.69) is located near the center of the PU.1 occupancy site (Fig. 4b,g).

Multiple lines of evidence support the regulatory effect of rs12517864. We reanalyzed ATAC-seq and histone mark ChIP-seq data for LCLs^36, 37^ and found that rs12517864 is significantly associated with each of these molecular phenotypes that overlap the PU.1 binding site, suggesting that the variant, if causal, affects gene regulation (Fig. 4f, h). Furthermore, the variant falls within a fragment that physically interacts only with the *ZNF608* promoter in primary B cells according to promoter-capture Hi-C (PCHi-C) data^38^, supporting the model that rs12517864 directly regulates *ZNF608* (Fig. 4g).

Surprisingly, initial inspection of *ZNF608* eQTL signals in LCLs^39^ seemed contradictory because the lead variant for this eQTL (rs2028854) is located elsewhere, 200 kb upstream of the *ZNF608* promoter, and is not strongly associated with lymphocyte count^21^ (*p* = 0.04) (Fig. 4c, g). We therefore examined the possibility of multiple independent *ZNF608* eQTL signals in LCLs by performing conditional analysis on the lead variant, as well as fine-mapping using SuSiE^40^, which can detect multiple signals. Once conditioned on the lead eQTL SNP rs2028854, association of rs12517864 to *ZNF608* expression became much stronger (*p* = 6.98×10^−7^) (Fig. 4c). Moreover, the fine-mapping analysis identified two independent credible sets for *ZNF608* eQTL signal, one of which contained rs12517864 as the variant with the highest posterior inclusion probability (PIP = 0.07), demonstrating that this variant is likely to be causally associated with *ZNF608* expression level (Fig. 4d).

Since only one of the two independent *ZNF608* eQTL signals in LCLs is associated with lymphocyte count, we hypothesized that even though both SNPs are significant eQTLs in LCLs, only rs12517864 (*i.e.*, the secondary eQTL signal), and not rs2028854 (*i.e.*, the primary eQTL signal), modulates *ZNF608* expression in the causal cell type. Analysis of RNA-seq data for various blood cells^28^ revealed that *ZNF608* is highly expressed in common lymphoid progenitors and B cells (Fig. 4i). Inspection of eQTL data for B cells in the eQTL Catalogue^41, 42^ showed that only rs12517864, and not rs2028854 (*p* = 0.25), is significantly associated with *ZNF608* expression (*p* = 4.39×10^−5^) (Fig. 4e). Although we cannot unambiguously conclude that B cells are the causal cell type, rs12517864 is likely the only variant increased lymphocyte count through increased *ZNF608* expression (Fig. 4h and Extended Data Fig. 3a).

### Blood cell trait-associated PU.1 motif-altering variants show regulatory effects in reporter assays

To verify that the nominated PU.1 motif-altering variants are indeed regulatory variants, we inspected massively parallel reporter assay (MPRA) studies data^43, 44^, which measured the regulatory effects of two such variants: (1) rs5827412, a PU.1 motif-altering short deletion associated with monocyte percentage affects expression levels of *LRRC25*, a gene previously shown to be necessary for granulocyte differentiation^45^, in monocytes (Fig. 5); and (2) rs3808619, a PU.1 motif-altering SNP at the promoter of *ZC2HC1A*, a functionally uncharacterized gene, as the regulatory causal variant for association with lower lymphocyte count (Fig. 6).

*LRRC25*, also called monocyte and plasmacytoid-activated protein (MAPA), is a gene shown to impair differentiation of granulocytes, which share lineages with monocytes, if knocked down or knocked out^45^. At this locus, we found that the PU.1 bQTL signal showed significant colocalization with monocyte count and percentage, neutrophil count and percentage, and white blood cell count association signals^8, 21^ (Fig. 5b and Extended Data Fig. 4a). The corresponding PU.1 binding site contains a short deletion rs5827412 that lowers the PU.1 motif score and is associated with reduced PU.1 binding, as well as chromatin accessibility, active histone mark levels, and *LRRC25* expression^36, 37, 39^ (Fig. 5a and Extended Data Fig. 4b). This deletion significantly reduced regulatory activity in a reporter assay^44^ (two-sided *t*-test *p* = 6.9×10^−5^) (Fig. 5f); data from another study suggested concordant direction of effect despite not being statistically significant^43^ (negative binomial regression *p* = 0.26) (Fig. 5c). Next, we analyzed available ATAC-seq data from *SPI1*, the gene encoding PU.1, knockout pro-B cell lines (RS4;11) to verify whether PU.1 is likely to be the trans factor for the regulatory variant^46^, and determined that *SPI1* knockout resulted in significantly reduced chromatin accessibility at sites of PU.1 occupancy genome-wide^47^ (chi square test *p* < 1×10^−300^) (Supplementary Fig. 3). Indeed, the activity of the regulatory element that contains rs5827412 is likely dependent on PU.1 binding as *SPI1* knockout cell lines showed reduced chromatin accessibility at this region (DESeq2 adjusted *p* = 8.73×10^−5^) (Fig. 5d). RNA-seq data for 13 blood cell types^28^ indicates that *LRRC25* is specifically expressed in monocytes at a much higher level than in other blood cell types and is sharply upregulated as progenitor cells differentiate to monocytes (Fig. 5e and Extended Data Fig. 4c). Consistent with the variant’s strongest effect on monocyte percentage (*p* = 1.3×10^−96^) and monocyte-specific expression of *LRRC25*, we found that rs5827412 is also significantly associated with reduced *LRRC25* expression in monocytes^16^ (*p* = 3.78×10^−22^) (Fig. 5f) and is in a regulatory element that is accessible throughout monocyte differentiation (Fig. 5g). Altogether, our results provide strong support for rs5827412 reducing *LRRC25* gene expression levels in monocytes and decreasing monocyte percentage while increasing neutrophil percentage.

The *ZC2HC1A* locus, which is primarily associated with lymphocyte count and percentage^21^ (Fig. 6b and Extended Data Fig. 5a,b), represents a challenging locus for fine-mapping. Here, 44 variants comprise the 95% credible set (*i.e.*, a minimal set of putative causal variants), based on a UK Biobank fine-mapping study^48^ (Fig. 6c). Among the candidate causal variants at the *ZC2HC1A* locus, rs3808619 is the only PU.1 motif-altering variant found within the associated PU.1 binding site at the *ZC2HC1A* promoter; rs3808619 increases the strength of a PU.1 motif, resulting in a higher affinity DNA binding site (Fig. 6a). Of multiple tagging variants in this locus that were tested for reporter activity (59 variants in Abell et al.^43^ and 30 variants in Tewhey et al.^44^), only rs3808619 showed a significantly increased reporter activity that is concordant in direction with that of the variant’s associations to elevated chromatin accessibility, active histone mark levels, and *ZC2HC1A* expression in LCLs^36, 37, 39^ (Fig. 6d,e,f). Finally, as for rs5827412, we detected significantly reduced chromatin accessibility levels at the *ZC2HC1A* promoter in *SPI1* knockout cell lines^46^ (DESeq2 adjusted *p* = 1.76×10^−13^), supporting the likely role of PU.1 at this promoter (Fig. 6g). rs3808619 is also associated with multiple sclerosis^49^ (*p* = 1.1×10^−9^) (Extended Data Fig. 5c,d), suggesting it plays a multifactorial role in IMDs. Our results suggest that a direct consequence of rs3808619, which is associated with lower lymphocyte count, is likely *ZC2HC1A* upregulation (Supplementary Note).

## Discussion

Our results with PU.1 binding and blood cell trait GWAS data demonstrate the utility of TF bQTL data in identifying which of many variants in LD are the likely causal regulatory variants underlying GWAS trait associations, as the presence of motif-altering variants suggests that they directly affect binding of the corresponding TF (Fig. 2c). Incorporating PU.1 bQTLs in our colocalization analysis conferred two key advantages: 1) identification of trait-associated regulatory elements and 2) identification of putatively causal PU.1 motif-altering variants. Together, they highlight a likely transcriptional regulatory mechanism underlying the trait association. In contrast, eQTL colocalization cannot assist fine-mapping in this way because there is no prior expectation that a specific noncoding region regulates the associated gene and that a regulatory variant would alter a certain TF binding site.

For instance, at the *ZNF608* locus, pinpointing the putative causal variant and associated regulatory element would have been difficult without PU.1 bQTLs, especially because there is another stronger eQTL signal, which did not colocalize with the lymphocyte count association for *ZNF608* in LCLs (Fig. 4). Such a situation may partially explain the observation that many significant eQTL signals failed to colocalize with the GWAS associations using existing colocalization methods^13^; however, this locus was the only such example in our study. Nevertheless, this example motivates applying TF bQTL colocalization to isolate independent eQTL signals, generating eQTL data in trait-relevant cell types^50^, and applying colocalization methods that allow multiple causal variants to eQTLs^51^, if accurate LD matrices or individual genotypes are available for both traits, which is often not the case for GWAS data.

A prior study that performed colocalization analysis of PU.1 bQTLs in neutrophils and immune diseases GWAS found that the majority (>50%) of colocalized variants altered the binding site motifs of other TFs^18^; in contrast, we found that the majority (87%) of the colocalized blood cell trait GWAS loci had a variant that altered a PU.1 motif (Fig. 2c), even though only a minority (34%) of all PU.1 bQTLs, colocalized or not, did overall (Fig. 1c). The increased proportion of PU.1 motif-altering variants present in this study may be due to PU.1’s central role in blood cell traits^26^ and highlights the increased likelihood that PU.1 binding is mediating the genetic effects on blood cell traits.

We observed that only a minority of the tested GWAS loci (69 / 367) showed significant colocalization. This is not surprising because we selected candidate loci solely based on the marginal association to PU.1 binding and blood cell traits^13^, without filtering for high LD between the two lead variants^13^ to ‘cast a wide net’ for discovery. This observation is a testament to the importance of performing colocalization analysis to distinguish loci with a single causal variant for the two phenotypes (here, PU.1 binding and a particular blood cell trait) from those with distinct variants responsible for the different phenotypes. Furthermore, even though PU.1 bQTLs were enriched for blood cell traits association (Fig. 2a), they explain only a subset of all associated loci, likely indicating that other TFs are mediating genetic effects at other associated loci.

We offer guidelines for broad application of colocalization analysis with TF bQTLs. First, high-quality ChIP-grade antibodies^52^ or, alternatively, cell lines in which the TF has been epitope-tagged, are essential. Second, TFs for bQTL analysis, as well as the cell type for the ChIP experiments, must be selected to be relevant to the trait or disease of interest. The feasibility of our analysis relied on the relevance of PU.1, a known hematopoietic master regulator, and LCLs, a model of mature B cells, to specific blood cell traits, such as lymphocyte count and monocyte count. Future studies will need to validate the regulatory functions of the variants in the relevant primary cell types. Third, sufficiently large sample sizes for both GWAS and TF bQTL are necessary for discovery, as colocalization can return false negative results due to limited statistical power^53^; although the sample size of 49 for the PU.1 bQTL data led to 69 robustly colocalized loci, we anticipate that a larger sample size could increase the power to detect weaker colocalization signals.

Future studies could use TF bQTL data in colocalization analysis to elucidate the ever-increasing number of trait-associated loci^1^. Where TFs important for a trait are known, TF bQTLs identified in the relevant cell type(s) could mediate a subset of trait associations, shedding light on putative causal variants, as well as the pathogenic mechanisms. Such colocalization analysis with TF bQTL data uniquely provides a path to pinpointing causal regulatory elements and variants, and thus a smaller set of mechanistic hypotheses to test experimentally to verify the underlying causes of the disease.

## Methods

### PU.1 ChIP-seq data processing

We downloaded PU.1 ChIP-seq fastq files from EMBL-EBI ArrayExpress under accession E-MTAB-3657^22^ (*n*=45) and E-MTAB-1884^19^ (*n*=4). The list of samples is provided in Supplementary Table 1. We mapped the reads to the hg19 reference genome supplemented with the Epstein-Barr virus (EBV) using Bowtie 2^54^. In order to eliminate reference allele bias in read mapping, we applied WASP^55^ to filter reads that mapped to a different position when variants were added, and used GSNAP^56^, which is a SNP-tolerant read alignment method, to remap filtered out reads.

PU.1 ChIP-seq peaks were called using MACS2^57^. For equal representation, we subsampled 5 million reads from each sample and performed peak calling on the aggregate alignment file. To account for the size of the merged read set, we downloaded 8 available control ChIP-seq samples in GM12878 from ENCODE (File ID: ENCFF032WUR, ENCFF426WJH, ENCFF508HCX, ENCFF537DAJ, ENCFF812HUT, ENCFF837IOW, ENCFF849LYY, ENCFF892TNJ). To define 200 bp sequences occupied by PU.1, we took the summits and extended them 100 bp in each direction. In total, there were 78720 peaks.

### PU.1 binding quantitative trait loci

First, we quantified the PU.1 binding levels at identified occupancy sites. We counted the number of reads overlapping each 200 bp peak using featureCounts^58^. The read counts were normalized for library size using trimmed mean of M-values^59^ and further normalized to follow a standard normal distribution across the samples, using quantile normalization. Finally, in order to eliminate the effect of variables, such as batch, gender, and ancestry, we used PEER^60^ to residualize the phenotype values, correcting for batch (*i.e.*, which publication), sex, and 3 genotype principal components, as well as 10 PEER factors.

Second, we obtained the genotypes of the LCL samples from the 1000 Genomes Project data^29^. 4 out of 49 samples only had microarray genotype data from Illumina Omni2.5 chips, and these genotypes were phased and imputed using the European samples of the 1000 Genomes project phase 3 data^29^ on the Michigan Imputation Server^61^. Genotypes of all samples were converted to biallelic form and aggregated. Afterwards, variants with minor allele frequency less than 5% were removed from the PU.1 binding quantitative trait loci analysis.

Finally, we tested for genetic associations to PU.1 binding levels using the phenotype matrix and the genotype data. We utilized QTLtools^62^ to approximate linear regression efficiently while also correcting for multiple hypotheses tested with permutations and false discovery rate estimation. For each PU.1 occupancy site, variants within 100 kb were included in the QTL analysis. In the end, there were 1497 significant PU.1 bQTLs.

### UK Biobank blood cell trait GWAS summary statistics

We downloaded 28 blood cell trait GWAS summary statistics from UK Biobank^21^ for the colocalization analysis. The authors performed a linear mixed model-based regression analysis on 452,264 White British individuals using rank-normalized phenotypes. The 28 blood cell traits are listed in Supplementary Table 3. One limitation of these summary statistics is that the authors used the Haplotype Reference Consortium imputation panel, which only included SNPs by design, for imputation^63^ (Supplementary Note). Thus, short deletions like rs5827412 were missing in these summary statistics. For Figure 5, we verified that the variant is associated with decreased monocyte percentage and increased neutrophil percentage in summary statistics from another analysis of the UK Biobank data^8^, and utilized these data for visualization.

### Fold Enrichment of GWAS signal in PU.1 bQTLs

We first generated 250 sets of null variants matched with the significant PU.1 bQTL lead variants for allele frequency, number of tagging SNPs (LD *r*^2^ > 0.5), and distance to the closest transcription start site (TSS), using SNPsnap^64^. 250 sets of null variants were successfully generated for 1292 of the PU.1 bQTL lead variants, so we restricted the downstream analysis within them. Using the distribution of number of variants tagging (*r*^2^ > 0.8) trait-associated lead variants as the background, we computed the fold enrichment of the number of PU.1 bQTLs tagging those variants. The empirical *p* values are derived for each blood cell trait by counting how many sets had SNPs tagging (*r*^2^ > 0.8) trait-associated variants more than or equal to the number of PU.1 bQTLs tagging them and dividing by 251. The *p* values were adjusted using *qvalue* package in R. For non-blood traits, lead SNPs from GWAS of type 2 diabetes^65^ and height^66^ were used.

### Position weight matrix and gkm-SVM PU.1 motif models

To initially scan for the position of PU.1 motif sequences within occupancy sites, we used PWMScan^67^. With a PU.1 (SPI1) motif position weight matrix (PWM) selected within the tool (CISBP: M6119_1) we scanned for the motif (*p* < 10^−5^) within PU.1 occupancy sites, which resulted in a total of 30812 instances. To determine the relative location of PU.1 motifs within the PU.1 occupancy sites, we subtracted the start or end position of the motif from the center position of the 200 bp PU.1 peak, depending on the strand (Extended Data Fig. 1a).

Afterwards, we trained a PU.1 motif model using gkm-SVM^31^, as a more sophisticated counterpart to PWM. We used the 200 bp sequences detected to be PU.1 occupancy sites for positive sequences in the training set. We left out PU.1 occupancy sites with a variant overlapping PU.1 motifs identified using PWMs (*i.e.*, one of the alleles with log-likelihood score > 8) from the training set so that the model effectively captures the motif sequences and excludes potentially causal PU.1 bQTLs. We generated negative sequences using the ‘genNullSeqs’ function in the gkmSVM R package. Then, we trained the model using default parameters with LS-GKM^31^, which is a faster implementation from the developers. Throughout the study, we defined PU.1 motif-altering variants as those where one of the alleles shows a gkm-SVM score greater than 0 for a 30 bp sequence centered at the variant, and the variant induces a non-zero change.

### Colocalization analysis using JLIM and Coloc

We selected 1621 PU.1-trait pairs at loci where the significant PU.1 bQTLs also show at least one blood cell trait association at *p* < 10^−6^ to perform colocalization. For JLIM^13^, we used the default parameters. *p* values were derived by permuting the PU.1 binding level matrix. For Coloc^12^, we used the prior parameters *p*1=10^−4^, *p*2=10^−4^, and *p*12=10^−6^, which is more conservative than the default, and ran Coloc on the summary statistics. For both analyses, we considered variants within a 200 kb window around the GWAS lead variant. We used a significance threshold of p < 0.01172 (FDR < 5%) for JLIM and posterior probability of colocalization (PP(Colocalization)) > 0.5. The FDR cutoff for JLIM was determined by the equation:

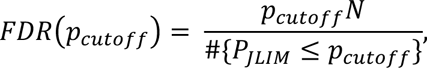

where *pcutoff* is the *p* value cutoff, *N* is the number of PU.1-trait loci tested, and *PJLIM* is the JLIM *p* value.

### Chromatin accessibility, histone mark, and expression QTLs in LCLs

ATAC-seq^36^ (*n*=100), histone mark ChIP-seq (*n*=158^13^ and *n*=2^34^, respectively), and RNA-seq^39^ (*n*=373) data were downloaded from European Nucleotide Archive (ERP110508), EMBL-EBI ArrayExpress (E-MTAB-3657 and E-GEUV-1), respectively. ATAC-seq data were only available as bam files, so we used bamtofastq command from bedtools^68^ to extract reads. We processed ATAC-seq and histone mark ChIP-seq read data similarly to PU.1 ChIP-seq data (*i.e.*, alignment, duplicate removal, peak calling, quantification, and then PEER^60^ normalization). The processed gene expression matrix derived from RNA-seq was downloaded directly.

We obtained the genotypes of the LCL samples from the 1000 Genomes Project data. We imputed 9 out of 100, 9 out of 160, and 15 out of 373 samples, respectively, from available microarray data to the 1000 Genomes Project phase 3 data^29^ on the Michigan Imputation Server^61^. Common variants (MAF > 5%) from the merged genotypes and the prepared phenotype matrices were used to test genetic associations to the corresponding molecular phenotypes with QTLtools^62^.

### Chromatin accessibility and gene expression levels across blood cell types

ATAC-seq and RNA-seq data from multiple blood cell types throughout hematopoiesis were downloaded from GEO series GSE74912 and GSE74246, respectively^28^. We aligned ATAC-seq read data to the hg19 reference genome, and merged data from each cell type for visualization. The genome tracks in Fig. 5 were generated with fold enrichment over average genome coverage to account for library size differences. We downloaded the count matrix for RNA-seq and converted them to counts per million for comparison across cell types.

### MPRA data analysis

We downloaded MPRA analysis tables from the two studies^43, 44^. We extracted statistics for rs5827412 and rs3808619, which were the only two putative causal PU.1 motif-altering variants at colocalized loci with MPRA data. For rs3808619, we also extracted the statistics for the other 29 and 58 variants tagging rs3808619 from Tewhey et al. and Abell et al., respectively. From Tewhey et al. data, we referred to the combined LCL analysis statistics, and from Abell et al. data, we referred to the allele effect statistics to measure the regulatory effects of variants.

### Differential accessibility analysis in *SPI1* knockout RS4;11 lines

ATAC-seq data from wild type and *SPI1* knockout RS4;11 cell lines were downloaded from EMBL-EBI ArrayExpress under accession E-MTAB-8676^46^. We aligned the reads using Bowtie2^54^ and removed duplicate alignments using scripts from WASP^55^. Then, we pooled the three replicates per genotype to call accessible regions using MACS2^57^ with *q* < 0.05 cutoff, and the two sets of accessible regions were merged using bedtools^68^. After counting the number of reads from each region using featureCount^58^, we tested for differential accessibility using DESeq2^69^. PU.1 ChIP-seq and input DNA data from unstimulated RS4;11 cell lines were downloaded from GEO series GSE71616^47^. After alignment using Bowtie2^54^ and duplicate removal^55^, we called peaks using MACS2^57^. Accessible regions were stratified by whether they intersect identified PU.1 occupancy sites. The significance of observing reduced accessibility in *SPI1* knockout lines was tested using a chi square test.

## Data availability

Processed data for generating the figures presented in the manuscript are available at https://github.com/BulykLab/PU1-colocalization-manuscript. PU.1 and Histone mark ChIP-seq data are available from EMBL-EBI ArrayExpress under accession <u>E-MTAB-3657</u> and <u>E-MTAB-1884</u>. ATAC-seq data in LCLs are available under European Nucleotide Archive accession <u>ERP110508</u>. Processed RNA-seq data in LCLs are available under EMBL-EBI ArrayExpress under accession <u>E-GEUV-1</u>. The 1000 Genomes Project Phase 3 genotype data are available at ftp://ftp.1000genomes.ebi.ac.uk/vol1/ftp/release/20130502. UK Biobank blood cell traits GWAS data from Canela-Xandri et al.^21^ are available at http://geneatlas.roslin.ed.ac.uk/, and those from Vuckovic et al.^8^ are available at ftp://ftp.sanger.ac.uk/pub/project/humgen/summary_statistics/UKBB_blood_cell_traits. Monocyte eQTL data from BLUEPRINT^16^ are available at http://blueprint-dev.bioinfo.cnio.es/WP10/qtls. Naïve B cell eQTL data from the eQTL Catalogue^41^ are available at ftp://ftp.ebi.ac.uk/pub/databases/spot/eQTL/sumstats/Schmiedel_2018/ge/Schmiedel_2018_ge_monocyte. <u>all.tsv.gz</u>. ATAC-seq data from control and *SPI1* knockout RS4;11 cell lines are available under EMBL-EBI ArrayExpress accession <u>E-MTAB-8676</u>, and PU.1 ChIP-seq data from RS4;11 cell line are available under GEO series accession <u>GSE71616</u>. Fine-mapping results for blood cell trait GWAS are available at https://github.com/bloodcellgwas/manuscript_code/tree/master/data/finemap_bedfiles/ukbb_v2 and https://www.finucanelab.org/data.

## Code availability

Codes for generating the figures are available at https://github.com/BulykLab/PU1-colocalization-manuscript. We trained a PU.1 motif gkm-SVM model using LS-GKM (https://github.com/Dongwon-Lee/lsgkm). We performed genotype imputation using the Michigan Imputation Server (https://imputationserver.sph.umich.edu/). We processed genotype data using BCFtools (https://samtools.github.io/bcftools/bcftools) and PLINK (https://www.cog-genomics.org/plink2/). We estimated hidden factors for QTL analyses using PEER (https://github.com/PMBio/peer). We generated sets of null variants for PU.1 bQTL enrichment analysis using SNPsnap (https://data.broadinstitute.org/mpg/snpsnap/). We performed colocalization analysis using JLIM (https://github.com/cotsapaslab/jlim) and Coloc (https://chr1swallace.github.io/coloc/). The Fuji plot (Fig. 3b) was made using code from https://github.com/mkanai/fujiplot. We adapted codes from LocusCompareR (https://github.com/boxiangliu/locuscomparer) to create association plots. We performed fine-mapping analysis for *ZNF608* eQTL in LCLs using SuSiE (https://stephenslab.github.io/susieR/).

## Acknowledgments

We thank members of the Bulyk lab, Vijay Sankaran, Shamil Sunyaev, Alexander Gusev, and members of the Raychaudhuri lab, including, but not limited to, Soumya Raychaudhuri, Kazuyoshi Ishigaki, Saori Sakaue, Tiffany Amariuta, Yang Luo, and Samira Asgari for helpful discussion and Shubham Khetan and Shamil Sunyaev for critical reading of the manuscript. This work was funded by a grant from the Brigham Research Institute Fund to Sustain Research Excellence and NIH grant R01 HG010501.

## Author Contributions

R.J. and M.L.B. conceived and designed the research project. R.J. performed all analyses and prepared the figures. M.L.B. supervised the research. R.J. and M.L.B. wrote the manuscript.

## Ethics Declarations

The authors declare no competing interests.

---

**Extended Data Fig. 1:**
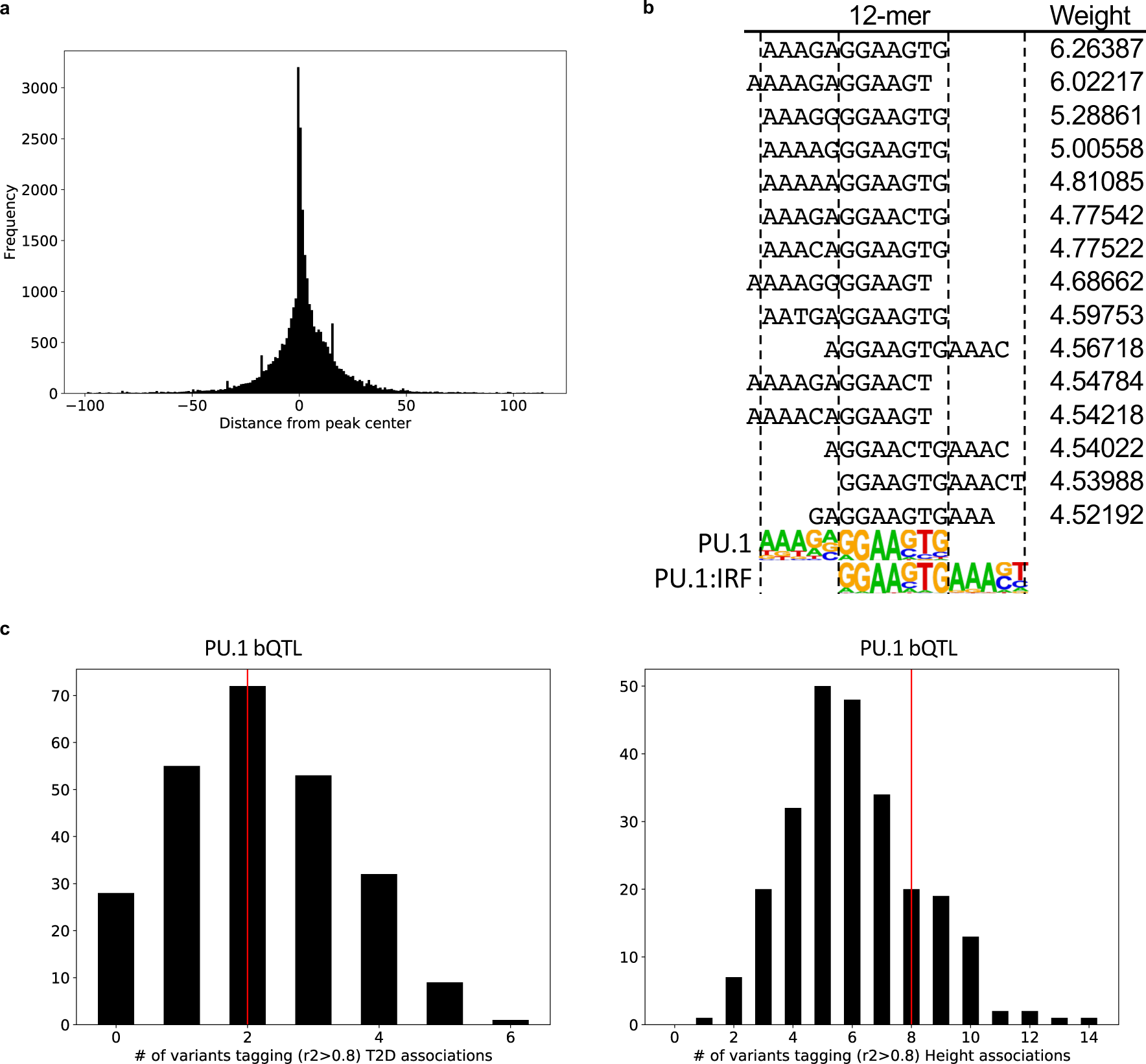
Properties of PU.1 binding sites and bQTLs. (**a**) Position of PU.1 motifs at PU.1 binding sites. The bp distance is measured from the center of a 200 bp PU.1 ChIP-seq peak. (**b**) 12-mers with the highest (top 15) gkm-SVM weights aligned to PU.1 motif and PU.1:IRF composite motif. (**c**) Lack of enrichment in PU.1 bQTL lead variants tagging (LD *r*^2^ > 0.8) type 2 diabetes (T2D) and height GWAS associations. The histogram shows the number of variants tagging GWAS associations for each of 250 sets of null variants. The red lines indicate the number of PU.1 bQTL lead variants tagging GWAS associations.

**Extended Data Fig. 2:**
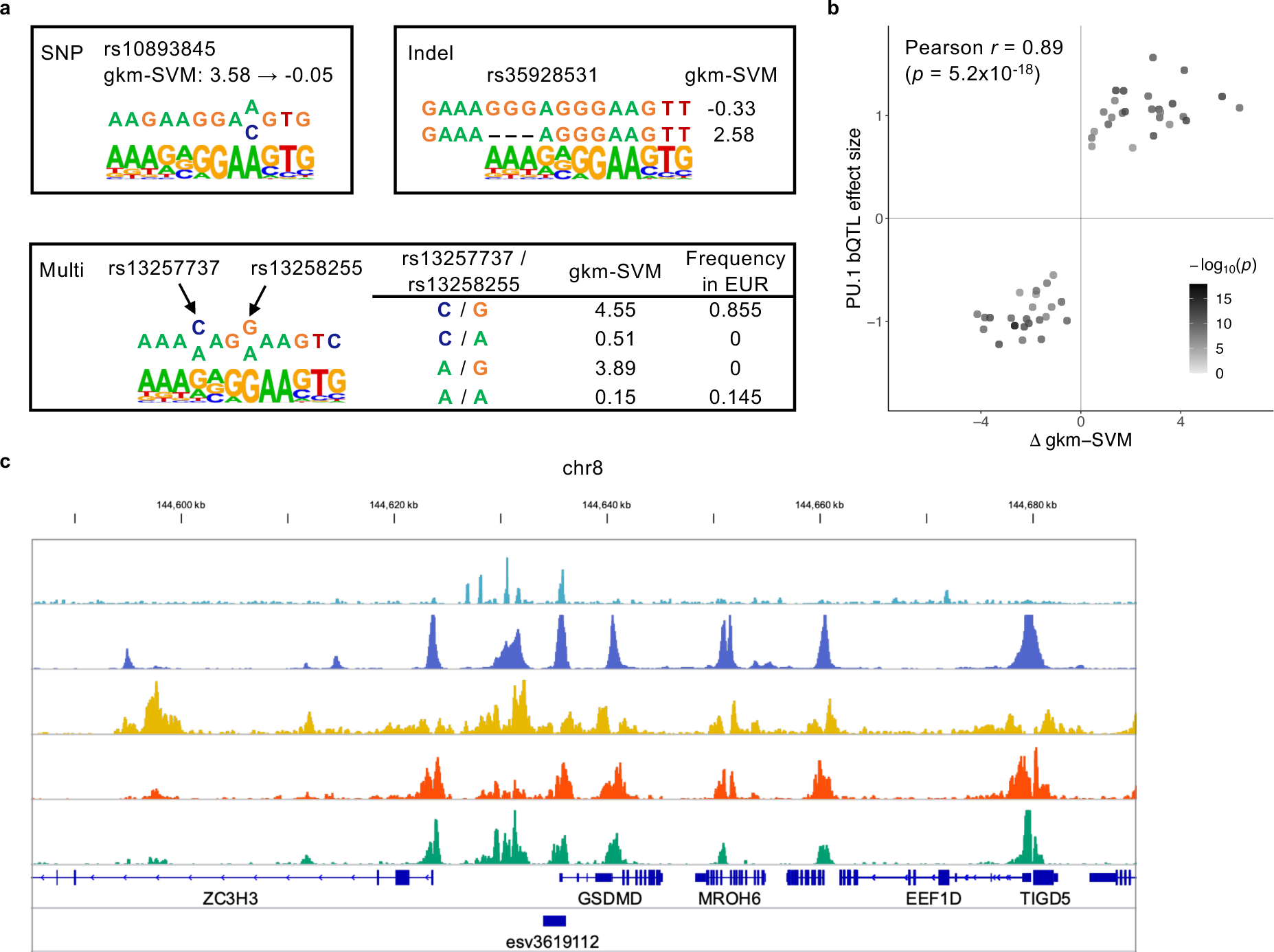
Examples of variants affecting PU.1 binding. (**a**) Examples of PU.1 motif-altering variants. Categorization of the variants correspond to Fig. 2b. EUR: European ancestry population in the 1000 Genomes Project. (**b**) Comparison of changes in motif score (Δ gkm-SVM) and estimated bQTL effect sizes of PU.1 motif-altering variants (SNPs and indels) at 49 colocalized loci. (**c**) An example of a copy number variation (esv3619112) affecting a PU.1 binding site.

**Extended Data Fig. 3:**
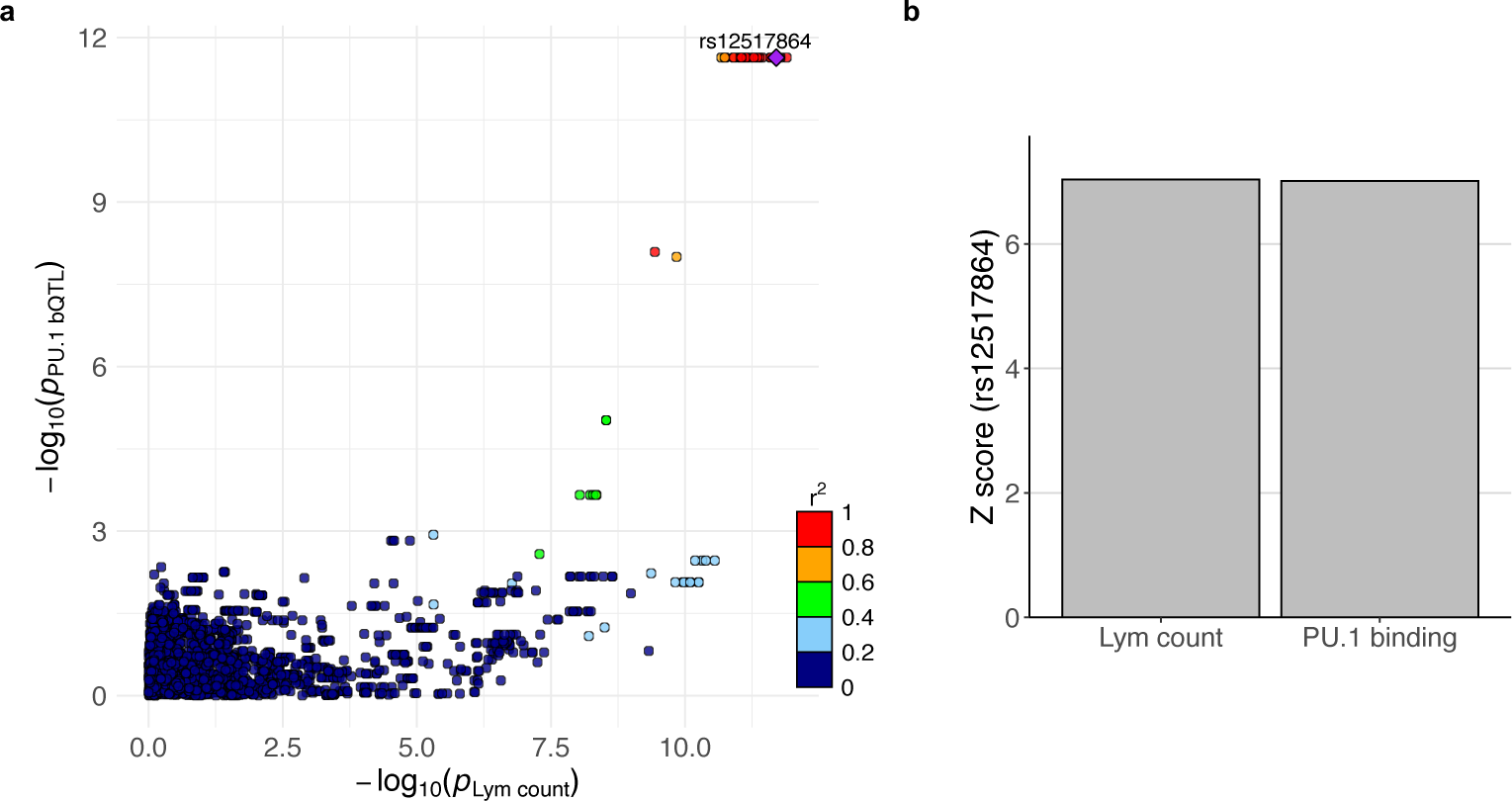
Colocalization of PU.1 bQTL and lymphocyte count association signals at *ZNF608* locus. (**a**) Merged association plot for PU.1 bQTL and lymphocyte count association signals. Points are colored by LD *r*^2^ with respect to rs12517864, which is labeled with a purple diamond. (**b**) Z scores of rs12517864 for lymphocyte count and PU.1 bQTL association.

**Extended Data Fig. 4:**
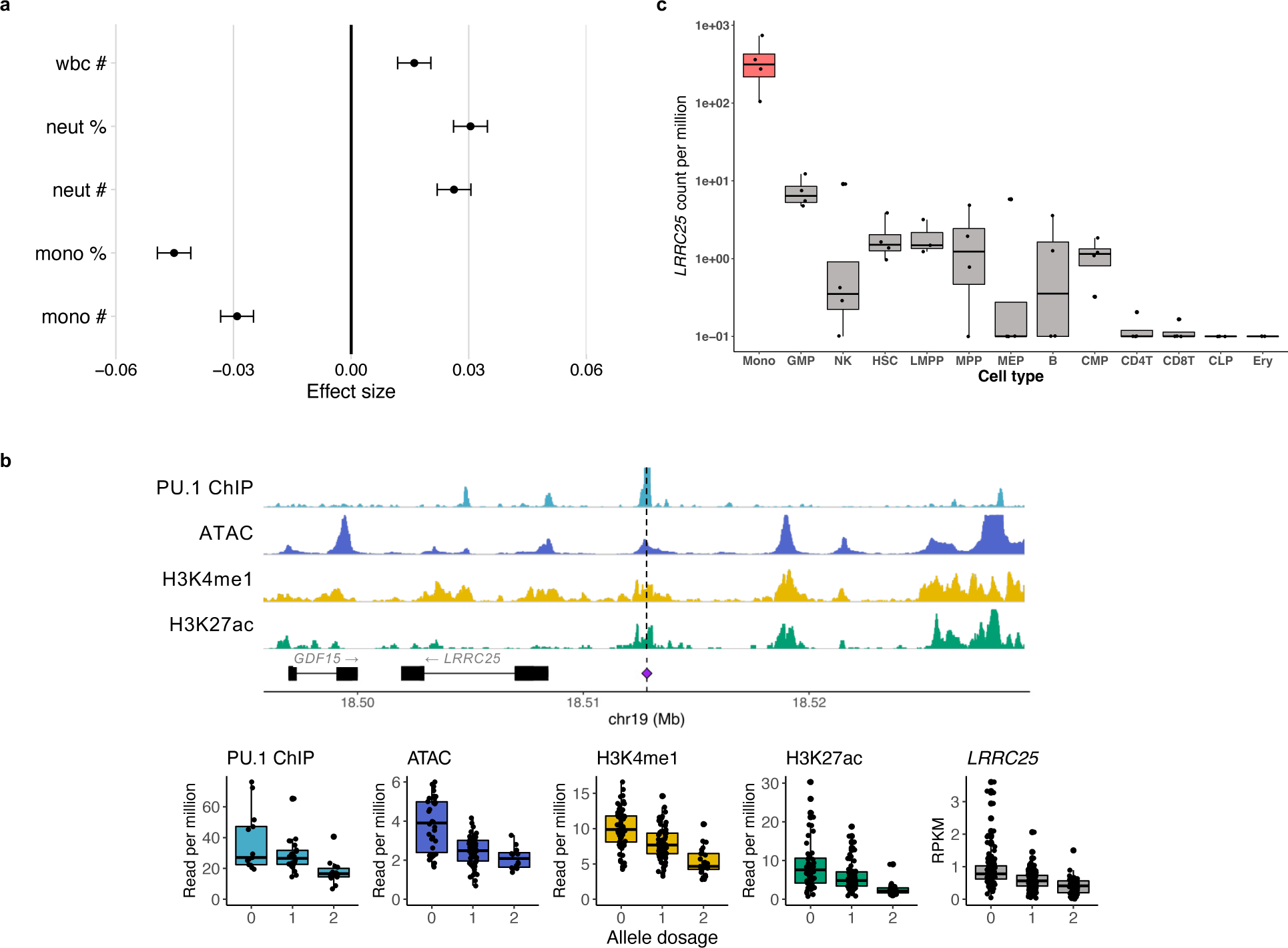
Effects of PU.1 motif-altering deletion rs5827412. (**a**) GWAS effect size estimates for rs5827412 on 5 blood cell traits. The error bars indicate 95% confidence interval. Abbreviations of blood cell traits are described in Supplementary Table 2. (**b-c**) Boxplots are formatted as in Fig 4. (**b**) Regulatory QTL effects of rs5827412. (top) Genome tracks show PU.1 ChIP-seq, ATAC-seq, and H3K4me1 and H3K27ac ChIP-seq data from LCLs, respectively. (bottom) 4 phenotype values in read per million for each genome track and reads per kilobase million for *LRRC25* expression levels. Allele dosage corresponds to the deletion allele. (**c**) *LRRC25* expression level across 13 blood cell types. Monocyte is colored red. Cell types abbreviated as in Supplementary Fig. 1.

**Extended Data Fig. 5:**
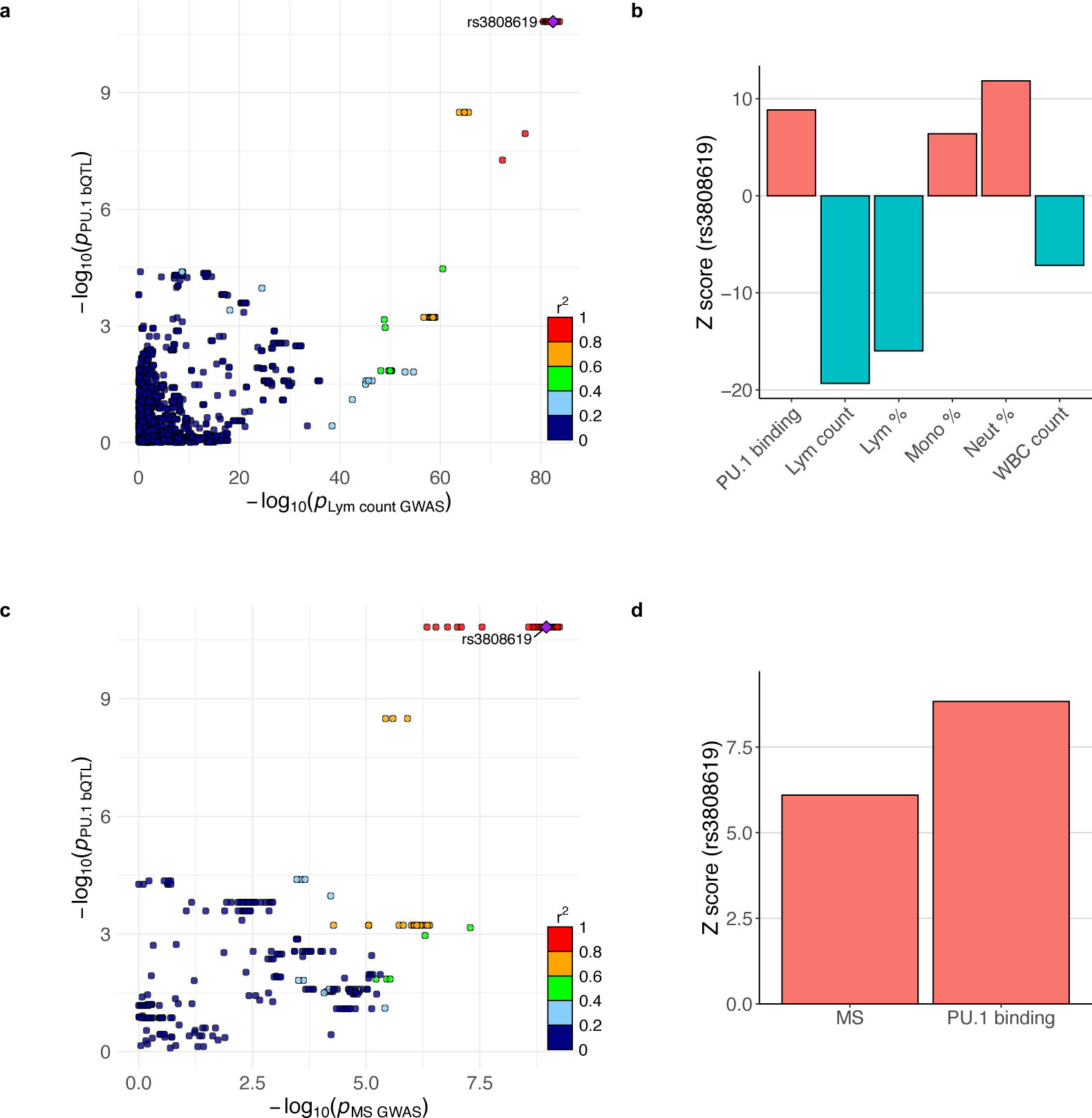
Colocalization of PU.1 bQTL and multiple sclerosis association signals at *ZC2HC1A* locus. (**a,c**) Points are colored by LD *r*^2^ in the 1000 Genomes Project European population, with respect to rs3808619, which is labeled with a purple diamond. (**a**) Merged association plot for PU.1 bQTL and lymphocyte count association signals. (**b**) Z scores of rs3808619 for PU.1 bQTL and 5 blood cell traits association. (**c**) Merged association plot for PU.1 bQTL and multiple sclerosis (MS) association signals. (**d**) Z scores of rs3808619 for MS and PU.1 bQTL association.

## Notes

### Competing Interest Statement

The authors have declared no competing interest.

## References

1. Claussnitzer, M. et al. A brief history of human disease genetics. Nature 577, 179–189 (2020).

2. Claussnitzer, M., Dankel, S. N., Kim, K.-H., Hauner, H. & Kellis, M. FTO obesity variant circuitry and adipocyte browning in humans. New England Journal of Medicine vol. 6 895–907 (2015).

3. Nasser, J. et al. Genome-wide enhancer maps link risk variants to disease genes. Nature 593, 238– 243 (2021).

4. International Common Disease Alliance. International Common Disease Alliance White Paper v1.0. https://www.icda.bio/ (2020).

5. Visscher, P. M. et al. 10 Years of GWAS Discovery: Biology, Function, and Translation. Am. J. Hum. Genet. 101, 5–22 (2017).

6. Amariuta, T. et al. Improving the trans-ancestry portability of polygenic risk scores by prioritizing variants in predicted cell-type-specific regulatory elements. Nat. Genet. 52, 1346–1354 (2020).

7. Weissbrod, O. et al. Leveraging fine-mapping and multipopulation training data to improve cross-population polygenic risk scores. Nat. Genet. 54, 450–458 (2022).

8. Vuckovic, D. et al. The Polygenic and Monogenic Basis of Blood Traits and Diseases. Cell 182, 1214–1231 (2020).

9. Amariuta, T. et al. IMPACT: Genomic Annotation of Cell-State-Specific Regulatory Elements Inferred from the Epigenome of Bound Transcription Factors. Am. J. Hum. Genet. 104, 879–895 (2019).

10. van de Geijn, B. et al. Annotations capturing cell type-specific TF binding explain a large fraction of disease heritability. Hum. Mol. Genet. 29, 1057–1067 (2020).

11. Ulirsch, J. C. et al. Interrogation of human hematopoiesis at single-cell and single-variant resolution. Nat. Genet. 51, 683–693 (2019).

12. Giambartolomei, C. et al. Bayesian test for colocalisation between pairs of genetic association studies using summary statistics. PLoS Genet. 10, e1004383 (2014).

13. Chun, S. et al. Limited statistical evidence for shared genetic effects of eQTLs and autoimmune-disease-associated loci in three major immune-cell types. Nat. Genet. 49, 600–605 (2017).

14. GTEx Consortium. The GTEx Consortium atlas of genetic regulatory effects across human tissues. Science 369, 1318–1330 (2020).

15. Barbeira, A. N. et al. Exploiting the GTEx resources to decipher the mechanisms at GWAS loci. Genome Biol. 22, 49 (2021).

16. Chen, L. et al. Genetic Drivers of Epigenetic and Transcriptional Variation in Human Immune Cells. Cell 167, 1398–1414.e24 (2016).

17. Liu, B., Gloudemans, M. J., Rao, A. S., Ingelsson, E. & Montgomery, S. B. Abundant associations with gene expression complicate GWAS follow-up. Nat. Genet. 51, 768–769 (2019).

18. Watt, S. et al. Genetic perturbation of PU.1 binding and chromatin looping at neutrophil enhancers associates with autoimmune disease. Nat. Commun. 12, 1–12 (2021).

19. Kilpinen, H. et al. Coordinated effects of sequence variation on DNA binding, chromatin structure, and transcription. Science 342, 744–747 (2013).

20. Deplancke, B., Alpern, D. & Gardeux, V. The Genetics of Transcription Factor DNA Binding Variation. Cell 166, 538–554 (2016).

21. Canela-Xandri, O., Rawlik, K. & Tenesa, A. An atlas of genetic associations in UK Biobank. Nat. Genet. 50, 1593–1599 (2018).

22. Waszak, S. M. et al. Population Variation and Genetic Control of Modular Chromatin Architecture in Humans. Cell 162, 1039–1050 (2015).

23. Guan, W.-J. et al. Clinical Characteristics of Coronavirus Disease 2019 in China. N. Engl. J. Med. 382, 1708–1720 (2020).

24. Terpos, E. et al. Hematological findings and complications of COVID-19. Am. J. Hematol. 95, 834–847 (2020).

25. Wang, S., Sheng, Y., Tu, J. & Zhang, L. Association between peripheral lymphocyte count and the mortality risk of COVID-19 inpatients. BMC Pulm. Med. 21, 55 (2021).

26. Fisher, R. C. & Scott, E. W. Role of PU.1 in hematopoiesis. Stem Cells 16, 25–37 (1998).

27. Rothenberg, E. V, Hosokawa, H. & Ungerbäck, J. Mechanisms of Action of Hematopoietic Transcription Factor PU.1 in Initiation of T-Cell Development. Front. Immunol. 10, 228 (2019).

28. Corces, M. R. et al. Lineage-specific and single-cell chromatin accessibility charts human hematopoiesis and leukemia evolution. Nat. Genet. 48, 1193–1203 (2016).

29. Auton, A. et al. A global reference for human genetic variation. Nature 526, 68–74 (2015).

30. Ghandi, M., Lee, D., Mohammad-Noori, M. & Beer, M. A. Enhanced Regulatory Sequence Prediction Using Gapped k-mer Features. PLoS Comput. Biol. 10, (2014).

31. Lee, D. et al. A method to predict the impact of regulatory variants from DNA sequence. Nat. Genet. 47, 955–961 (2015).

32. Escalante, C. R. et al. Crystal structure of PU.1/IRF-4/DNA ternary complex. Mol. Cell 10, 1097–1105 (2002).

33. Yan, J. et al. Systematic analysis of binding of transcription factors to noncoding variants. Nature 591, 147–151 (2021).

34. Hormozdiari, F. et al. Colocalization of GWAS and eQTL Signals Detects Target Genes. Am. J. Hum. Genet. 99, 1245–1260 (2016).

35. Krysiak, K. et al. Recurrent somatic mutations affecting B-cell receptor signaling pathway genes in follicular lymphoma. Blood 129, 473–483 (2017).

36. Kumasaka, N., Knights, A. J. & Gaffney, D. J. High-resolution genetic mapping of putative causal interactions between regions of open chromatin. Nat. Genet. 51, 128–137 (2019).

37. Delaneau, O. et al. Chromatin three-dimensional interactions mediate genetic effects on gene expression. Science 364, (2019).

38. Javierre, B. M. et al. Lineage-Specific Genome Architecture Links Enhancers and Non-coding Disease Variants to Target Gene Promoters. Cell 167, 1369–1384 (2016).

39. Lappalainen, T. et al. Transcriptome and genome sequencing uncovers functional variation in humans. Nature 501, 506–511 (2013).

40. Wang, G., Sarkar, A., Carbonetto, P. & Stephens, M. A simple new approach to variable selection in regression, with application to genetic fine mapping. J. R. Stat. Soc. Ser. B Stat. Methodol. 82, 1273–1300 (2020).

41. Kerimov, N. et al. A compendium of uniformly processed human gene expression and splicing quantitative trait loci. Nat. Genet. 53, 1290–1299 (2021).

42. Schmiedel, B. J. et al. Impact of Genetic Polymorphisms on Human Immune Cell Gene Expression. Cell 175, 1701–1715 (2018).

43. Abell, N. S. et al. Multiple causal variants underlie genetic associations in humans. Science 375, 1247–1254 (2022).

44. Tewhey, R. et al. Direct identification of hundreds of expression-modulating variants using a multiplexed reporter assay. Cell 165, 1519–1529 (2016).

45. Liu, W. et al. LRRC25 plays a key role in all-trans retinoic acid-induced granulocytic differentiation as a novel potential leukocyte differentiation antigen. Protein Cell 9, 785–798 (2018).

46. Coz, C. Le et al. Constrained chromatin accessibility in PU.1-mutated agammaglobulinemia patients. J. Exp. Med. 218, (2021).

47. Wu, J. N. et al. Functionally distinct patterns of nucleosome remodeling at enhancers in glucocorticoid-treated acute lymphoblastic leukemia. Epigenetics Chromatin 8, 53 (2015).

48. Kanai, M. et al. Insights from complex trait fine-mapping across diverse populations. Preprint at https://www.medrxiv.org/content/10.1101/2021.09.03.21262975v1 (2021).

49. International Multiple Sclerosis Genetics Consortium (IMSGC) et al. Analysis of immune-related loci identifies 48 new susceptibility variants for multiple sclerosis. Nat. Genet. 45, 1353–60 (2013).

50. Umans, B. D., Battle, A. & Gilad, Y. Where Are the Disease-Associated eQTLs? Trends Genet. 37, 109–124 (2021).

51. Wallace, C. A more accurate method for colocalisation analysis allowing for multiple causal variants. PLoS Genet. 17, 1–11 (2021).

52. Baker, M. Reproducibility crisis: Blame it on the antibodies. Nature 521, 274–6 (2015).

53. Hukku, A. et al. Probabilistic Colocalization of Genetic Variants from Complex and Molecular Traits: Promise and Limitations. Am. J. Hum. Genet. 108, 25–35 (2020).

54. Langmead, B. & Salzberg, S. L. Fast gapped-read alignment with Bowtie 2. Nat. Methods 9, 357– 359 (2012).

55. Van De Geijn, B., Mcvicker, G., Gilad, Y. & Pritchard, J. K. WASP: Allele-specific software for robust molecular quantitative trait locus discovery. Nat. Methods 12, 1061–1063 (2015).

56. Wu, T. D. & Nacu, S. Fast and SNP-tolerant detection of complex variants and splicing in short reads. Bioinformatics 26, 873–881 (2010).

57. Zhang, Y. et al. Model-based analysis of ChIP-Seq (MACS). Genome Biol. 9, R137 (2008).

58. Liao, Y., Smyth, G. K. & Shi, W. FeatureCounts: An efficient general purpose program for assigning sequence reads to genomic features. Bioinformatics 30, 923–930 (2014).

59. Robinson, M. D. & Oshlack, A. A scaling normalization method for differential expression analysis of RNA-seq data. Genome Biol. (2010).

60. Stegle, O., Parts, L., Piipari, M., Winn, J. & Durbin, R. Using probabilistic estimation of expression residuals (PEER) to obtain increased power and interpretability of gene expression analyses. Nat. Protoc. 7, 500–507 (2012).

61. Das, S. et al. Next-generation genotype imputation service and methods. Nat. Genet. 48, 1284– 1287 (2016).

62. Delaneau, O. et al. A complete tool set for molecular QTL discovery and analysis. Nat. Commun. 8, 15452 (2017).

63. McCarthy, S. et al. A reference panel of 64,976 haplotypes for genotype imputation. Nat. Genet. 48, 1279–1283 (2016).

64. Pers, T. H., Timshel, P. & Hirschhorn, J. N. SNPsnap: a Web-based tool for identification and annotation of matched SNPs. Bioinformatics 31, 418–20 (2015).

65. Mahajan, A. et al. Fine-mapping type 2 diabetes loci to single-variant resolution using high-density imputation and islet-specific epigenome maps. Nat. Genet. 50, 1505–1513 (2018).

66. Wood, A. R. et al. Defining the role of common variation in the genomic and biological architecture of adult human height. Nat. Genet. 46, 1173–1186 (2014).

67. Ambrosini, G., Groux, R. & Bucher, P. PWMScan: a fast tool for scanning entire genomes with a position-specific weight matrix. Bioinformatics 34, 2483–2484 (2018).

68. Quinlan, A. R. & Hall, I. M. BEDTools: a flexible suite of utilities for comparing genomic features. Bioinformatics 26, 841–2 (2010).

69. Love, M. I., Huber, W. & Anders, S. Moderated estimation of fold change and dispersion for RNA-seq data with DESeq2. Genome Biol. 15, 1–21 (2014).

