## Supplemental information for "Colocalization of blood cell traits GWAS associations and variation in PU.1 genomic occupancy prioritizes causal noncoding regulatory variants"

### Supplementary Note

#### Note about discordant results from JLIM and Coloc

Although we didn't aim to rigorously investigate the differences between JLIM[1] and Coloc[2], we looked through the examples where the two methods showed discordant results (Fig. 2b, Supplementary Fig. 2). First, we visually inspected the association plots for some of the loci, where only Coloc showed significant colocalization. Here, we could not clearly determine whether they are false positives by Coloc or false negatives by JLIM (Supplementary Fig. 2a). It is possible that the LD structure is different enough between the GWAS cohort and the PU.1 bQTL samples to cause JLIM to fail to reject the null hypothesis. On the other hand, loci that only JLIM showed colocalization often had a large set of variants in LD (Supplementary Fig. 2b). This trend is likely due to JLIM's model specification, where JLIM statistics is higher if the lead variants for the two traits show high LD[1], even if the LD block includes more variants. In sum, some of the loci with discordant results can be false negatives, but we decided to focus on loci with significant colocalization from both methods.

#### Note about the two blood cell traits GWAS data

We utilized two blood cell traits GWAS data for this work[3,4]. They are both statistics for the UK Biobank data with notable differences. Canela-Xandri and colleagues analyzed data for 452,264 White British individuals, whereas Vuckovic and colleagues analyzed those from 408,112 individuals of British ancestry. They both applied linear mixed models. We incorporated Canela-Xandri et al. data for colocalization analyses because we expected greater statistical power due to larger sample sizes. However, Canela-Xandri and colleagues imputed the genotypes using the Haplotype Reference Consortium panel, which only includes SNPs and not indels, leading to SNP-only data. On the other hand, Vuckovic and colleagues imputed the genotypes using 1000 Genomes Project Phase 3[5] and UK10K[6] panel, which includes SNPs and short indels. Therefore, we used Vuckovic et al. data for plotting Figure 5, where a short deletion alters the PU.1 motif, and for determining credible set sizes based on their fine-mapping results.

#### Note about lymphocyte count association at *ZC2HC1A* locus

We pinpointed the PU.1 motif-altering SNP rs3808619 as the likely regulatory variant for colocalized PU.1 bQTL and lymphocyte count association at *ZC2HC1A* locus. Since the variant affects a PU.1 motif at its binding site at *ZC2HC1A* promoter, and the variant is significantly associated with increased *ZC2HC1A* expression, we hypothesized that the direct consequence of the variant is *ZC2HC1A* upregulation. As *ZC2HC1A* has no known function yet, we investigated this locus further. *IL7* gene is located downstream of *ZC2HC1A*, and a multi-ancestry blood cell trait GWAS study[7] demonstrated that a South Asian ancestry-specific missense mutation (rs2014122253) in *IL7* that increased IL-7 protein secretion in a heterologous cellular system was associated with increased lymphocyte count. rs2014122253 is extremely rare in the European population, so it is not in LD with rs3808619. Interestingly, in eQTLGen data, rs3808619 was significantly, but relatively weakly, associated ($p$=9.45x10$^{-14}$) with lower *IL7* expression[8] (this is compared to $p$=3.27x10$^{-310}$ for *ZC2HC1A*). Although our analysis with GEUVADIS European LCL samples[9] didn't show significant association ($p > 0.1$), eQTL Catalogue data[10] showed that rs3808619 is significantly associated with lower *IL7* expression in multi-ancestry GEUVADIS LCL eQTL analysis[9] ($p$ = 2.85x10$^{-9}$) and TwinsUK LCL eQTL analysis[11]

($p$=2.32x10$^{-10}$); only the latter analysis showed rs3808619 within the credible set of 41 variants. As Chen and colleagues showed that increased IL-7 secretion is associated with increased lymphocyte count[7], rs3808619's association with lower *IL7* expression and lower lymphocyte count is plausible. How rs3808619 increases regulatory activity by increasing affinity to PU.1 binding leading to increased *ZC2HC1A* expression potentially lowers *IL7* expression is yet unresolved.

### Reference


1. Chun, S. *et al.* Limited statistical evidence for shared genetic effects of eQTLs and autoimmune-disease-associated loci in three major immune-cell types. *Nat. Genet.* **49**, 600–605 (2017).
2. Giambartolomei, C. *et al.* Bayesian test for colocalisation between pairs of genetic association studies using summary statistics. *PLoS Genet.* **10**, e1004383 (2014).
3. Canela-Xandri, O., Rawlik, K. & Tenesa, A. An atlas of genetic associations in UK Biobank. *Nat. Genet.* **50**, 1593–1599 (2018).
4. Vuckovic, D. *et al.* The Polygenic and Monogenic Basis of Blood Traits and Diseases. *Cell* **182**, 1214–1231 (2020).
5. Auton, A. *et al.* A global reference for human genetic variation. *Nature* **526**, 68–74 (2015).
6. Walter, K. *et al.* The UK10K project identifies rare variants in health and disease. *Nature* **526**, 82–89 (2015).
7. Chen, M. *et al.* Trans-ethnic and Ancestry-Specific Blood-Cell Genetics in 746,667 Individuals from 5 Global Populations. *Cell* 1198–1213 (2020).
8. Võsa, U. *et al.* Large-scale cis- and trans-eQTL analyses identify thousands of genetic loci and polygenic scores that regulate blood gene expression. *Nat. Genet.* **53**, (2021).
9. Lappalainen, T. *et al.* Transcriptome and genome sequencing uncovers functional variation in humans. *Nature* **501**, 506–511 (2013).
10. Kerimov, N. *et al.* A compendium of uniformly processed human gene expression and splicing quantitative trait loci. *Nat. Genet.* **53**, 1290–1299 (2021).
11. Buil, A., Brown, A. A., Lappalainen, T. & Dermitzakis, E. T. Gene-gene and gene-environment interactions detected by transcriptome sequence analysis in twins. *Nat. Genet.* **47**, 88–91 (2015).

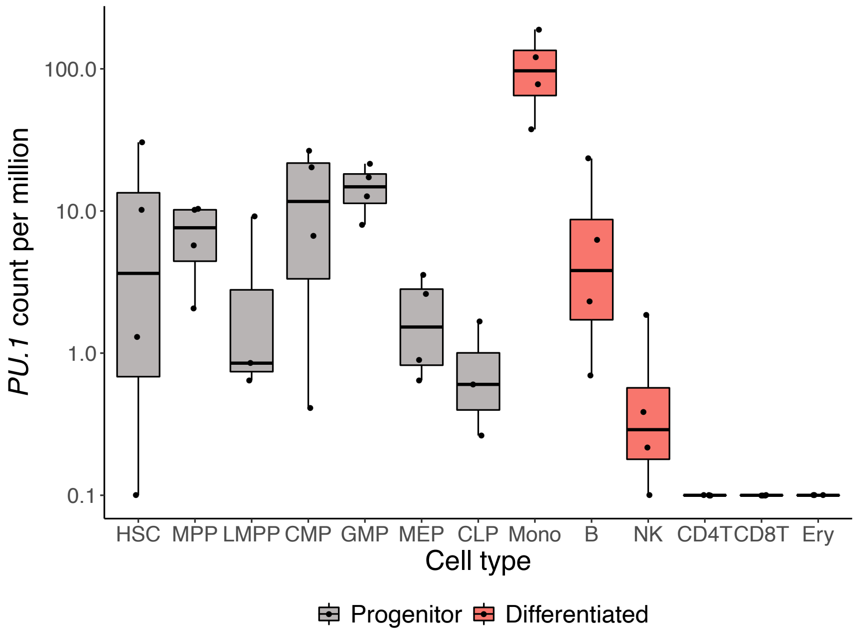


**Supplementary Fig. 1 | PU.1 expression across blood cell types.** Expression level is measured by bulk RNA-seq. Progenitor cell types (gray) and differentiated cell types (gray) are colored accordingly. HSC: hematopoietic stem cell, MPP: multipotent progenitor, LMPP: lymphoid-primed multipotent progenitor, GMP: granulocyte-monocyte progenitor, CMP: common myeloid progenitor, MEP: megakaryocyteCLP: common lymphoid progenitor, B: B cell, NK: natural killer cell, CD4T: $CD4^{+}$ T cell, CD8T: $CD8^{+}$ T cell, Ery: erythroid.

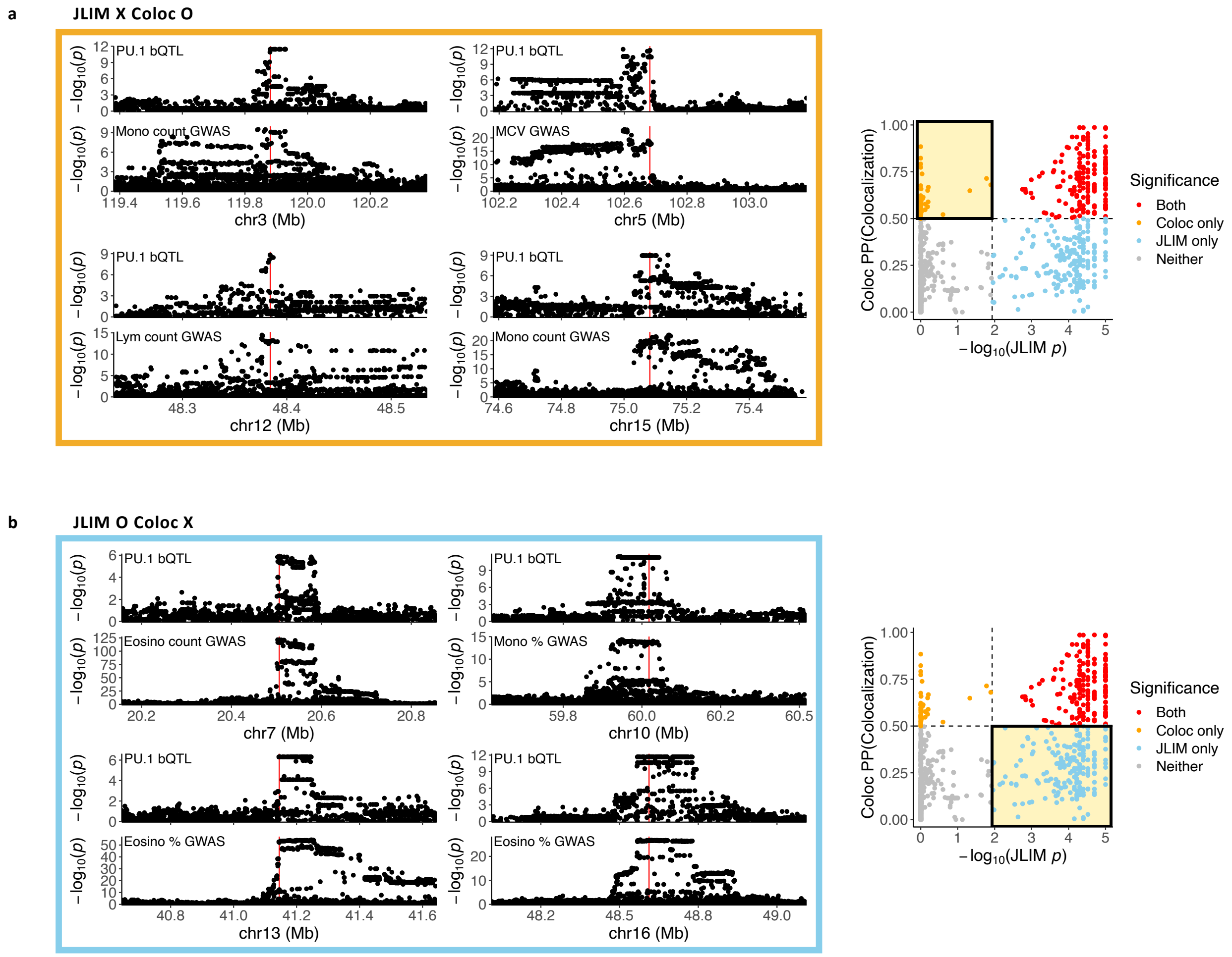


**Supplementary Fig. 2 | Examples of discordant colocalization results between JLIM and Coloc.** (**a**-**b**) (Left) Example association plots for PU.1 bQTL and various blood cell traits. (Right) Colocalization results (Fig. 2b) with yellow shading for the corresponding examples. (**a**) Loci with significant colocalization based on Coloc, but not JLIM. (**b**) Loci with significant colocalization based on JLIM, but not Coloc.

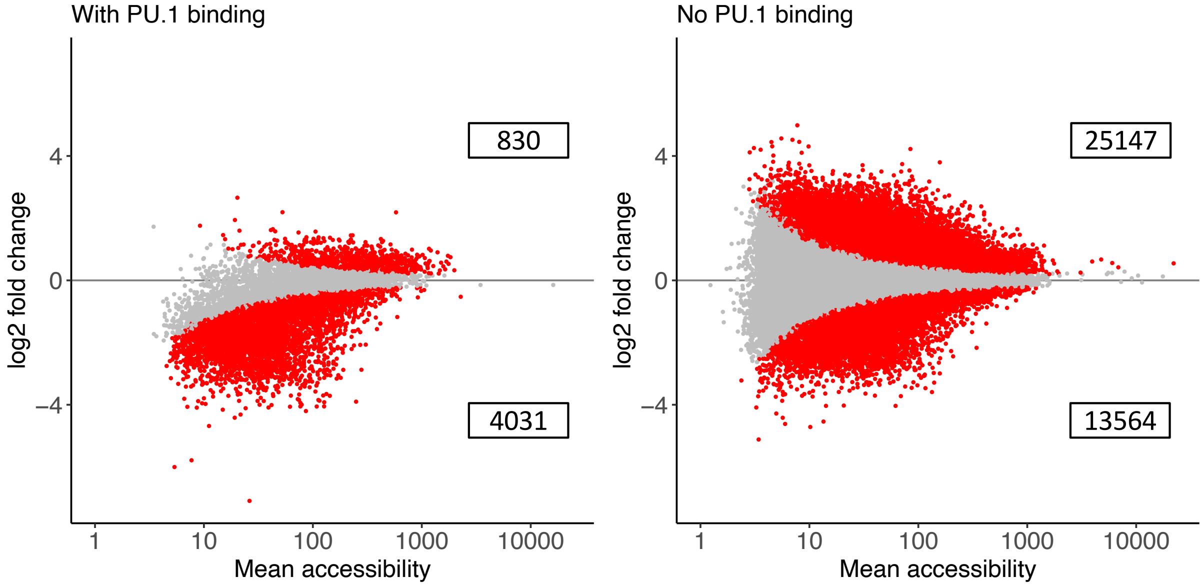


**Supplementary Fig. 3 | PU.1-dependent loss of chromatin accessibility.** Log2 fold change in chromatin accessibility in *SPI1*, the gene encoding PU.1, knock-out RS4;11 cell line for regions with PU.1 binding measured by ChIP-seq (left) and without PU.1 binding (right). Red points are accessible regions with significant gain or loss ($p_{adj} < 0.05$) of accessibility in knock-out mutants. Numbers in boxes represent the number of differentially accessible regions that either show increase or decrease, respectively, in accessibility in *SPI1* knock-outs.
